# Proteome dynamics during transition from exponential to stationary phase under aerobic and anaerobic conditions in yeast

**DOI:** 10.1101/2022.09.23.509138

**Authors:** Maxime den Ridder, Wiebeke van den Brandeler, Meryem Altiner, Pascale Daran-Lapujade, Martin Pabst

## Abstract

The yeast *Saccharomyces cerevisiae* is a widely used eukaryotic model organism and a promising cell factory for industry. However, despite decades of research, the regulation of its metabolism is not yet fully understood, and its complexity represents a major challenge for engineering and optimising biosynthetic routes. Recent studies have demonstrated the potential of resource and proteomic allocation data in enhancing models for metabolic processes. However, comprehensive and accurate proteome dynamics data that can be used for such approaches are still very limited. Therefore, we performed a quantitative proteome dynamics study to comprehensively cover the transition from exponential to stationary phase for both aerobically and anaerobically grown yeast cells. The combination of highly controlled reactor experiments, biological replicates and standardised sample preparation procedures ensured reproducibility and accuracy. Additionally, we selected the CEN.PK lineage for our experiments because of its relevance for both fundamental and applied research. Together with the prototrophic, standard haploid strain CEN.PK113-7D, we also investigated an engineered strain with genetic minimisation of the glycolytic pathway, resulting in the quantitative assessment of over 1700 proteins across 54 proteomes. These proteins account for nearly 40% of the overall yeast proteome and approximately 99% of the total protein biomass. The anaerobic cultures showed remarkably less proteome-level changes compared to the aerobic cultures, during transition from the exponential to the stationary phase as a consequence of the lack of the diauxic shift in the absence of oxygen. These results support the notion that anaerobically growing cells lack time and resources to adapt to changes in the environment. This proteome dynamics study constitutes an important step towards better understanding of the impact of glucose exhaustion and oxygen on the complex proteome allocation process in yeast. Finally, the established proteome dynamics data provide a valuable resource for the development of resource allocation models as well as for metabolic engineering efforts.

## INTRODUCTION

The yeast *Saccharomyces cerevisiae* is a widely used eukaryotic model organism and cell factory that represents a promising alternative to the fossil fuel-based production of chemicals. However, economic competitiveness is still a major hurdle for such cell factories. Constructing improved strains that realise high productivity and yield involves extensive genetic engineering to rewire native genomes that have been optimised for growth and survival over millions of years of evolution. Nevertheless, intensive research over the past decades have led to successful developments where yeast processes were brought to an industrial scale, such as for the production of the drug precursor artemisinic acid [1–4]. *In silico* approaches to reproduce and predict microbial metabolism have been simultaneously developed to assist metabolic engineering efforts [5]. However, the complexity of yeast metabolism limits the predictive power of these models. A promising approach to improve such models is to consider resource allocation and more particularly the cost of protein expression [6–10]. A prerequisite for this approach is the availability of comprehensive and accurate proteome dynamics data established under tightly controlled conditions. Unfortunately, such data are commonly not available and are difficult to obtain.

*S. cerevisiae* displays a remarkable metabolic flexibility, as it tunes its metabolism between full respiratory sugar dissimilation and alcoholic fermentation, with different degrees of respiro-fermentative metabolism as a function of environmental cues, substrate and oxygen supply. The well-known Crabtree effect results in partial repression of respiration and therefore in respiro-fermentative growth in the presence of excess sugar (e.g. glucose or galactose) even in aerobic conditions [11]. Conversely, the production of gluconeogenic substrates as ethanol or acetate leads to strict respiratory metabolism in aerobic settings. *S. cerevisiae* will fully ferment carbon sources in the absence of oxygen. However, respiratory and fermentative substrate dissimilation have a large impact on ATP yield, as full respiration of 1 mol of glucose results in 16 mol of ATP, while fermentation of the same amount of glucose only yields 2 mol of ATP [12]. The metabolic mode therefore strongly affects cellular resources, in particular their optimum allocation for growth and survival. To obtain a better insight into how *S. cerevisiae* responds to changes in substrate and oxygen supplies, we monitored its proteome employing tightly controlled bioreactors. Several yeast proteomics studies have already been performed over the past decades [13–21]. In this study, we monitored the dynamic proteome responses to substrate availability during all growth phases of yeast (exponential, diauxic and stationary phases) under both aerobic and anaerobic conditions [22]. Thus far, only little has been known regarding the proteome dynamics under anaerobic conditions, in particular during transition from the exponential to the stationary phase.

Considering eukaryotes, such as *S. cerevisiae*, genetic redundancy is another level of complexity for *in silico* design and experimental development of cell factories. Many genes, more particularly those involved in metabolism, have orthologues with similar functions [23], but often with a poorly understood physiological role. In view of minimal genomes, several studies have explored the requirement for these redundant genes and implemented top-down approaches to reduce genetic redundancy [24–26]. Such minimised genomes have the potential to facilitate the complete redesign and construction of entirely synthetic yeast genomes. Moreover, genetic minimisation of key metabolic pathways can facilitate the formulation and validation of mathematical models by eliminating isoenzymes with different regulatory and kinetic properties. Solis-Escalante *et al*. constructed a yeast strain in which the 26 genes encoding enzymes of the Embden–Meyerhof–Parnas pathway of glycolysis, the main pathway for sugar utilisation, were minimalised to a set of 13 genes in the minimal glycolysis (MG) strain (**Figure 1a**) [26]. While this genetically reduced strain appeared physiologically comparable to its parent strain (with the full set of glycolytic genes), the underlying proteome dynamics and potential protein level adjustments were not investigated. In this study, the engineered MG strain IMX372 and its parental *S. cerevisiae* CEN.PK113-7D were investigated together during transition from the exponential to the stationary phase in the presence or absence of oxygen. The temporal proteome dynamics across all growth phases were monitored from triplicate bioreactor cultures. Quantitative shotgun proteomic experiments were performed using 10-plex tandem mass tag (TMT) isobaric labelling. The use of tightly controlled reactor experiments in combination with robust sample preparation protocols allows the establishment of highly accurate quantitative data, which constitute valuable resources for *in silico* approaches, including metabolic engineering effort assistance. Furthermore, the established proteome dynamics data expand the current understanding of protein dynamics in yeast during carbon-limited growth under both aerobic and anaerobic conditions from the proliferation to the stationary phase. Finally, the comparison to the MG mutant quantified the impact of the loss of the minor glycolytic isoenzymes on the global proteome.

**FIGURE 1.**
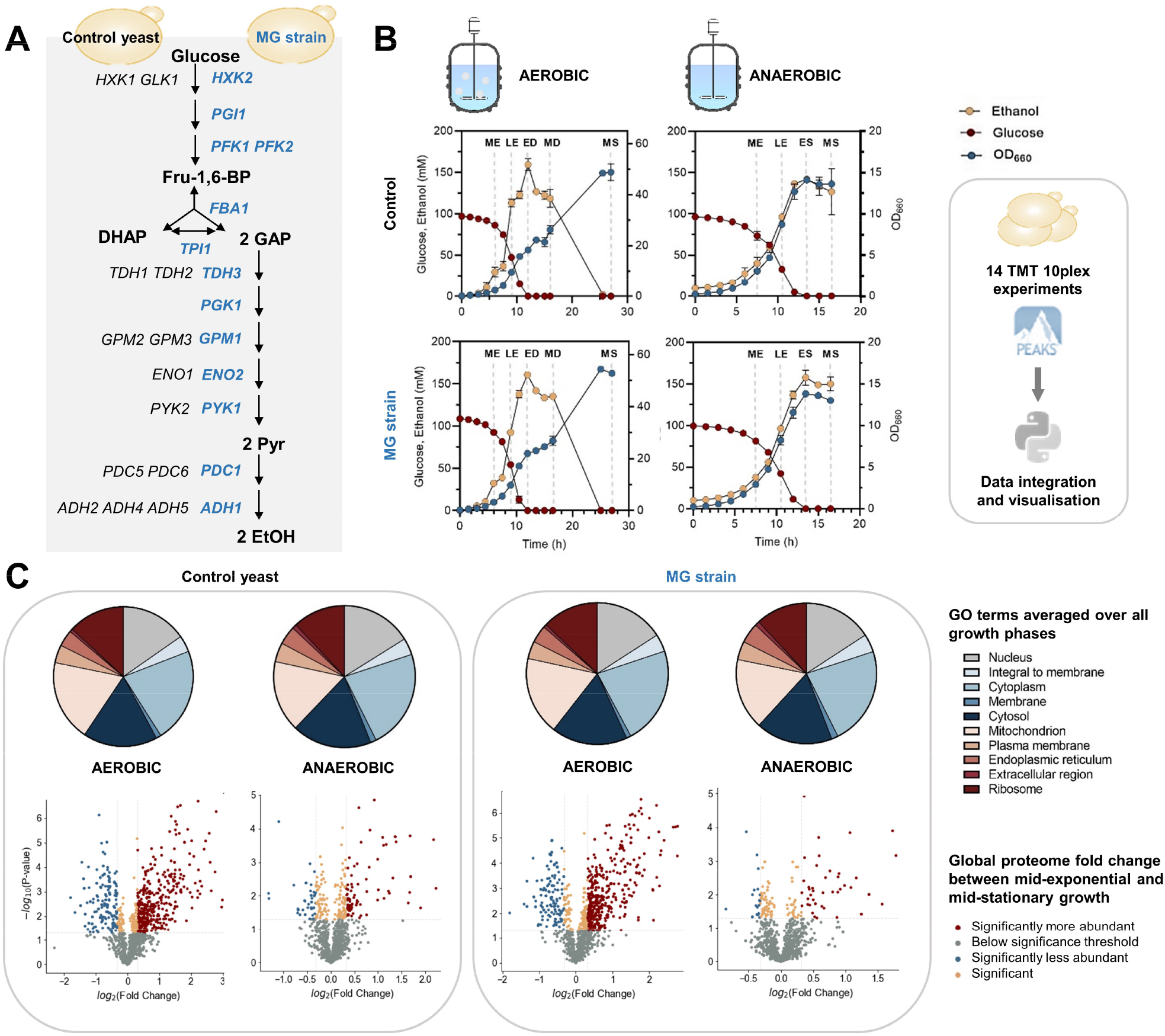
Yeast proteome dynamics study capturing the transition from proliferation to stationary phase under aerobic and anaerobic conditions. A) Schematic overview of glycolysis in the yeast control strain (CEN.PK113-7D, black) and the MG strain (IMX372, blue). The enzymes in blue are retained in MG yeast (*Adh3 is a mitochondrial protein). B) Yeast growth in aerobic and anaerobic cultures. Glucose (red) and ethanol (yellow) concentrations, and OD660 (blue, secondary y-axis) were measured during the different growth phases of aerobic and anaerobic batch cultures for the control yeast and the MG strain. The values shown are averages obtained from three biological replicates. Standard deviations are indicated by error bars. The dotted grey lines indicate time points at which samples were taken for proteome analysis. Proteome samples were taken from each biological replicate in the aerobic cultures after 6, 9, 12, 16.5 and 27 hours of growth, in the mid-exponential (ME), late-exponential (LE), early-diauxic (ED), mid-diauxic (MD) and (mid-) stationary (MS) growth phase, respectively. Furthermore, proteome samples were taken of the anaerobic cultures after 7.5, 10.5, 13.5 and 16.5 hours of growth, in the ME, LE, early-stationary (ES) and MS growth phase, respectively. Proteome samples were subjected to quantitative shotgun proteomics experiments, using 10-plex TMT isobaric labelling and a one-dimensional, 4-hour chromatographic separation. Database searching and quantitative analysis was performed using PEAKS X and using a tailor-made Python data processing pipeline. C) Annotation of yeast protein functions using Gene Ontology (GO) terms. Based on the classifications of GO annotation, the overall functions of the identified yeast proteins (with at least 2 unique peptides present) were categorized into cellular component, and displayed in pie chart format with absolute protein numbers (average of three biological replicates). The global proteome changes between the mid-exponential and mid-stationary phase under aerobic and anaerobic conditions in control and MG strain were visualised using volcano plots. The fold changes were normalized to the aerobic and anaerobic mid-exponential phases. The log2 of the abundance fold change between the two conditions was plotted against the significance (-log10p), using a p-value threshold of <0.05 and a fold change threshold of >1.25 (which corresponds to a log2 fold change threshold +/− 0.32). Significant changes of mid-stationary proteins were coloured by their direction of change (red if higher, blue if lower, or peach if similar to their mid-exponential equivalents). The total number of proteins with changes are listed in Table 1.

## EXPERIMENTAL SECTION

### Yeast strains and media

The MG yeast strain IMX372 (*MATa ura3-52 his3-1 leu2-3,112 MAL2-8c SUC2 glk1::SpHis5, hxk1::KlLEU2, tdh1::KlURA3, tdh2, gpm2, gpm3, eno1, pyk2, pdc5, pdc6, adh2, adh5, adh4*) and CEN.PK113-7D (*MATa MAL2-8C SUC2*) used in this study share the CEN.PK genetic background [26, 27]. Shake flask and batch cultures were grown in synthetic medium (SM) containing 5.0 g/L (NH_4_)SO_4_, 3.0 g/L KH_2_PO_4_, 0.5 g/L MgSO_4_·7H_2_O and 1 mL/L trace elements in demineralized water, set at pH 6. The medium was heat sterilized (120°C) and supplemented with 1 mL/L filter sterilized vitamin solution and 20 g/L heat sterilized (110 °C) glucose (SMG) [28]. The bioreactor medium was supplemented with 0.2 g/L antifoam Emulsion C (Sigma, St. Louise, USA) or with 0.2 g/L antifoam Pluronic PE 6100 (BASF, Ludwigshafen, Germany) for anaerobic and aerobic cultures, respectively. In case of anaerobic cultivations, the medium was also supplied with anaerobic growth factors, 10 mg/L ergosterol (Sigma-Aldrich, St. Louis, MO) and 420 mg/L Tween 80 (polyethylene glycol sorbate monooleate, Merck, Darmstadt, Germany) dissolved in ethanol. Frozen stocks of *S. cerevisiae* cultures were prepared by the addition of glycerol (30% v/v) in 1 mL aliquots for storage at −80 °C. **Bioreactor cultures.** Aerobic shake flask cultures were grown at 30°C in a Innova incubator shaker (New Brunswick™ Scientific, Edison, NJ, USA) at 200 rpm using 500 mL round-bottom shake flasks containing 100 mL medium. Triplicate aerobic batch cultures of control and MG yeast were performed in 2 L laboratory fermenters (Applikon, Schiedam, The Netherlands) with a 1.2 L working volume under aerobic and anaerobic conditions. SM-medium was used and maintained at pH 5 by the automatic addition of 2 M KOH. Mixing of the medium was performed with stirring at 800 rpm. Gas inflow was filter sterilized and compressed air (Linde Gas, Schiedam, The Netherlands) or nitrogen (<10 ppm oxygen, Linde Gas) was sparged via the bottom of the bioreactor at a rate of 500 mL/min, for aerobic and anaerobic cultures, respectively. Dissolved oxygen levels were measured with Clark electrodes (Mettler Toledo, Greifensee, Switzerland). The temperature of the fermenters was maintained at 30°C. The reactors were inoculated with exponentially growing shake flask cultures of *S. cerevisiae* strain IMX372 and CEN.PK113-7D to obtain an initial optical density (OD660) of approximately 0.2. Sampling for HPLC and OD660 measurements was done every 90 minutes. Proteome samples were taken at 6, 9, 12, 16.5, 27 and at 7.5, 10.5, 13.5, 16.5 hours in aerobic and anaerobic conditions, respectively. **Biomass, metabolites and gas measurements.** To monitor growth, OD660 measurements were performed on a JENWAY 7200 spectrophotometer (Cole-Parmer, Stone, UK). The biomass dry weight was determined in duplicate as described earlier [28]. For extracellular metabolite determinations, broth samples were centrifuged for 5 min at 13,000 g and the supernatant was collected for analysis with a Waters alliance 2695 HPLC (Waters Chromatography B.V., Etten-Leur, The Netherlands) with an Aminex HPX-87H ion exchange column (Biorad, Hercules, CA, USA). The HPLC was operated at 60°C and 5 mM of H2SO4 was used as mobile phase at a rate of 0.6 mL/min. Off-gas concentrations of CO2 and O2 were measured using an NGA 2000 analyser. Proteome samples (~3–5 mg dry weight) were taken from batch cultures. The samples were collected in multifold in trichloroacetic acid (TCA) (Merck Sigma, Cat. No. T0699) with a final concentration of 10%. Samples were centrifuged at 4000 g for 5 min at 4°C. Cell pellets were frozen at −80°C [29]. **Yeast cell lysis, protein extraction and proteolytic digestion.** Cell pellets of the aerobic and anaerobic cultures were resuspended in lysis buffer composed of 100 mM Triethylammonium bicarbonate (TEAB) containing 1% SDS and phosphatase/protease inhibitors. Yeast cells were lysed by glass bead milling by 10 cycles of 1 minute shaking alternated with 1 min rest on ice. Proteins were reduced by addition of 5 mM DTT and incubation for 1 hour at 37°C. Subsequently, the proteins were alkylated for 60 min at room temperature in the dark by addition of 50 mM acrylamide. Protein precipitation was performed by addition of four volumes of ice-cold acetone (−20°C), followed by 1 hour freezing at −20°C. The proteins were solubilized using 100 mM ammonium bicarbonate. Proteolytic digestion was performed by Trypsin (Promega, Madison, WI), 1:100 enzyme to protein ratio, and incubated at 37°C overnight. Solid phase extraction was performed with an Oasis HLB 96-well μElution plate (Waters, Milford, USA) to desalt the mixture. Eluates were dried using a SpeedVac vacuum concentrator at 50°C and frozen at −80°C. **Quantitative temporal proteome analysis.** Desalted peptides were reconstituted in 100 mM TEAB and TMT10-plex reagents (Thermo) were added from stocks dissolved in 100% anhydrous acetonitrile (ACN). Peptides were mixed with labels in a 1:8 ratio (12.5 μg to 100 μg) and incubated for 1 hour at 25°C and 400 rpm and the labelling reaction was stopped by addition of 5% hydroxylamine to a final concentration of 0.4%. Labelled peptides were then mixed in at approx. equal quantities. Two bridging samples were included in each TMT10-plex experiment to improve comparability between different experiments. The bridging sample was a mixture of the three biological replicates of MG yeast under aerobic conditions in the mid-stationary phase. Peptide solutions were diluted with water to obtain a final concentration of acetonitrile (ACN) lower than 5%. Solid phase extraction was performed to desalt the final peptide mixture. Desalted peptides were subsequently frozen at −80°C for 1 hour and dried by vacuum centrifugation. Peptides were finally resuspended in 3% ACN/0.01% TFA prior to MS-analysis to give an approximate concentration of 500 ng per μL. Samples were labelled as indicated in **SI table 2**. **Shotgun proteomic analysis.** An aliquot corresponding to approximately 1 μg protein digest was analysed using an one dimensional shot-gun proteomics approach [30]. Briefly, the samples were analysed using a nano-liquid-chromatography system consisting of an EASY nano-LC 1200, equipped with an Acclaim PepMap RSLC RP C18 separation column (50 μm x 150 mm, 2 μm, Cat. No. 164568), and a QE plus Orbitrap mass spectrometer (Thermo Fisher Scientific, Germany). The flow rate was maintained at 350 nL/min over a linear gradient from 5% to 25% solvent B over 180 min, then from 25% to 55% over 60 min, followed by back equilibration to starting conditions. Data were acquired from 5 to 240 min. Solvent A was H2O containing 0.1% formic acid (FA), and solvent B consisted of 80% ACN in H2O and 0.1% FA. The Orbitrap was operated in data-dependent acquisition (DDA) mode acquiring peptide signals from 385–1250 m/z at 70 K resolution in full MS mode with a maximum ion injection time (IT) of 75 ms and an automatic gain control (AGC) target of 3E6. The top 10 precursors were selected for MS/MS analysis and subjected to fragmentation using higher-energy collisional dissociation (HCD). MS/MS scans were acquired at 35 K resolution with AGC target of 1E5 and IT of 100 ms, 1.2 m/z isolation width and normalized collision energy (NCE) of 32. **Processing of mass spectrometric raw data.** Data were analysed against the proteome database from *Saccharomyces cerevisiae* (Uniprot, strain ATCC 204508 / S288C, Tax ID: 559292, July 2020) using PEAKS Studio X (Bioinformatics Solutions Inc., Waterloo, Canada) [31], allowing for 20 ppm parent ion and 0.02 m/z fragment ion mass error, 3 missed cleavages, acrylamide and TMT10 label as fixed and methionine oxidation and N/Q deamidation as variable modifications. Peptide spectrum matches were filtered against 1% false discovery rates (FDR) and identifications with ≥2 unique peptides. Changes in protein abundances between different time points using the TMT quantification option provided by the PEAKSQ software tool (Bioinformatics Solutions Inc., Canada). Auto normalization was used for quantitative analysis of the proteins, in which the global ratio was calculated from the total intensity of all labels in all quantifiable peptides. Quantitative analysis was performed using protein identifications containing at least 2 unique peptides, which peptide identifications were filtered against 1% FDR. The significance method for evaluating the observed abundance changes was set to ANOVA and the significance score was expressed as the −10xlog10(p), where p is the significance testing p-value. The p-value represents the likelihood that the observed change is caused by random chance. Results from PEAKSQ were exported to ‘proteins.csv’, containing the quantified proteins. **Pathway analysis, functional enrichment, and data visualisation.** Briefly, the exported ‘proteins.csv’ files from PEAKSQ, listing the quantified proteins for each experiment, were directly imported into the Python environment. Normalization between data was performed using a bridging sample. A function was further established that links Uniprot accession numbers and yeast genes (as obtained from https://www.uniprot.org/docs/yeast.txt, and which subsequently was used to annotate identified proteins from the experiments with correct gene names. The biological triplicates per condition (aerobic and anaerobic) and strain (control and MG) were treated separately. Furthermore, each biological replicate consisted of two additional technical replicates. To analyse the technical and biological replicates, clustermaps were made using a self-built Python function based on the clustermap function from the Seaborn package in Python [32], using the Euclidean distances metric and the average linkage method. Only proteins detected in all three biological replicates were used for the cluster analysis. The fold change of each protein in a specific condition was calculated relative to the bridging sample. The average fold changes of the technical replicates were subsequently used to determine the standard deviations of the biological replicates. The averages of the biological replicates were determined to obtain the four sub-datasets i) control aerobic, ii) MG aerobic iii) MG anaerobic and iv) control anaerobic. All graphs ultimately show the analyses of these biological-replicate averages and their corresponding standard deviations.

To study how protein abundances changed in individual cellular pathways, the obtained proteomics data were analysed using the KEGG (Kyoto Encyclopaedia of Genes and Genome) pathway database [33]. All the up-to-date KEGG pathways were retrieved with the constructed ‘KEGG_tool.py’ code. Here, the Bio.KEGG.REST module from the Biopython package in Python was used [34]. Thereby, the functions ‘kegg_list’ was used to list all pathways for *S. cerevisiae*, and ‘kegg_get’ to retrieve gene names that are assigned to a specific pathway. Since many pathways have an extensive list of members, the pathways in the central carbon metabolism (CCM) were reduced to the most important genes in order to enable meaningful visualisation in graphs. Using the above mentioned clustermap function, the protein fold changes of the CCM were plotted on a heatmap for each of the experiments, without any clustering. For better visualisation of the trends, the data were normalised to the mid-exponential (ME) phase. The same function was moreover used to display the average absolute intensity of every protein throughout the whole growth curve, using log10 and absolute scale, respectively. The significance of a difference in biological-replicate-average fold changes between two datasets was assessed by performing a two-sided two-sample unpaired t-test (also known as Welch’s t-test), using the ‘ttest_ind_from_stats’ function from the ‘SciPy.stats’ module in Python [35]. Global proteome changes between two experiments or phases were visualised in volcano plots, where the - 10log10(p) is plotted against the log2(fold change) between the two conditions. These plots were generated using the ‘gene_exp.volcano’ a modified version of the GeneExpression.volcano function from the ‘Bioinfokit.visuz’ module in Python [36]. This function enabled the division of the fold changes between two experiments into i) insignificant changes, ii) statistically significant changes (but not necessarily biologically significant), iii) statistically and likely biologically significant changes. For this study the statistical significance threshold was generally set to p <0.05. The (presumed) biological significance threshold was set to a log2 fold change threshold of +/− 0.32 (indicating a 1.25 absolute fold change).

A functional enrichment analysis using the STRING database was performed in order to determine whether specific GO-terms or KEGG-pathways are enriched under a particular condition [37]. For this, Python was used to programmatically accesses the STRING database via an API. This created a dictionary containing the up- and downregulated proteins, the species identifier (4932 for *S. cerevisiae*), the functional categories that should be assessed, the FDR threshold (<0.05 in this study), and an optional set of ‘background genes’ with as alternative background the whole species proteome. The function ‘backgroundgene_2_string’ retrieves the protein-specific string identifiers for the back-ground genes/proteins, which in this case were all proteins detected across the experiments. Estimation of the average protein content for the aerobic and anaerobic growth conditions using emPAI and PAI indices was performed according to Yasushi Ishihama et al., 2005 [38]. Circle graphs were made using the ‘surf’ function in Matlab, where circle areas represent the obtained emPAI values. **Data availability.** Mass spectrometric raw data have been deposited to the ProteomeXchange Consortium [39] via the PRIDE [40] partner repository and are publicly available under the project code PXD031412.

## RESULTS

### Proteome dynamics of laboratory control CEN.PK113-7D and MG yeast in aerobic and anaerobic batch bioreactor cultures

To optimise data reproducibility and reliability, we performed the batch cultures in bioreactors in which mixing, aeration and pH were tightly controlled. Independent triplicate cultures were conducted for the two investigated strains to further increase biological significance. Furthermore, we selected the prototrophic control strain *S. cerevisiae* CEN.PK113-7D – a popular lineage for biotechnology for which several omics datasets are already available – and the MG variant (IMX372) lacking glycolytic minor isoenzymes, for our study (**Figure 1a**). The batch cultures were sampled during all growth phases, ranging from the proliferation phase to growth arrest in the stationary phase (**Figure 1b**). Generally, the presence or absence of oxygen is known to strongly affect yeast physiology, which results in differences in metabolism and growth phases. During growth on glucose, aerobic cultures both respire and ferment, producing ethanol and other fermentation products. The growth on glucose is followed by a diauxic growth phase during which fermentation products are fully respired until the stationary phase. Conversely, *S. cerevisiae* fully ferments glucose and does not respire in the absence of oxygen. Dissimilation of fermentation products requires oxygen; therefore, anaerobic cultures directly switch from exponential growth on glucose to the stationary phase, without a diauxic phase. These physiological differences were also observed in the growth and metabolite profiles performed in our study (**SI Table 1 and Figure 1b**). Aerobic proteome dynamics were monitored in time with sampling at 6, 9, 12, 16.5 and 27 hours of growth, corresponding to the mid-exponential, late exponential, early diauxic, mid-diauxic and stationary growth phases, respectively (**Figure 1b**). To align the sampling time points to the physiology, we sampled the anaerobic cultures at 7.5, 10.5, 13.5 and 16.5 hours of growth, corresponding to the mid- and late exponential, early stationary and stationary growth phases, respectively (**Figure 1b**). After cell lysis and trypsin digestion, peptide samples of three biological replicates per condition were labelled using TMT10-plex reagents, mixed equally and subjected to a 4-hour (gradient) shotgun proteomics experiment (**SI Table 2**). On average, 1175 and 1106 proteins were quantified in the control yeast under aerobic and anaerobic conditions, respectively, with at least two unique peptides and 1% FDR. Similarly, 1131 and 1127 proteins were quantified confidently on average for the aerobic and anaerobic cultures of the MG strain, respectively. A total of 1734 proteins were quantified across all TMT experiments (**SI Table 3**), which is close to 40% of the total proteome considering the theoretical expression of approximately 4500 proteins at any time [41]. The protein amount was estimated using the emPAI index [38]. Furthermore, the total protein content was estimated by summing all emPAI values, thereby assuming for unidentified proteins the lowest observed emPAI value of our study. This indicated that more than 99% of the total protein biomass was captured in our study (**SI Figure 1 and SI Table 4**). The detected proteins were predominantly assigned to intracellular organelle functional GO categories, consisting of cytosolic, mitochondrial and ribosomal proteins (**Figure 1c and SI Table 5**), which could be explained by their high expression levels [19, 42]. A similar GO term assignment was found for both strains and conditions.

The crude protein quantification profiles were further analysed using a Python data processing pipeline to enable a tailored visualisation and interpretation of the large-scale data. To this end, the data from the 14 separate TMT experiments were compared for their temporal and conditional protein abundance changes in the control strain (CEN.PK113-7D) and MG yeast strain IMX372 under (an)aerobic conditions. Data were presented as fold changes for each protein in a specific condition relative to a bridging sample. The bridging (control) sample used in all TMT experiments was a mixture of the three biological replicates of the aerobic stationary-phase MG yeast to improve comparability between the experiments (**SI Table 2**).

To assess experimental reproducibility, we compared global proteomic data with cluster analysis data based on Euclidean distances using three biological replicates per strain and condition and two technical replicates per time point within each biological replicate. This demonstrated clustering of replicates of the different growth phases per strain and condition, confirming the reproducibility of the reactor experiments and proteomic analyses (**SI Figure 2**). The average protein abundance of the three biological replicates per condition and strain was further used for the interpretation of the proteome dynamics data.

### Effects of oxygen availability on the global proteome dynamics across the growth curve

To explore the impact of oxygen availability on the yeast proteome, we first focused on the growth phases with the most marked change in the global proteome between the aerobic and anaerobic cultures for the control strain CEN.PK113-7D (i.e. stationary and mid-exponential phases). While a similar number of proteins were quantified in the presence and absence of oxygen, the number of differently expressed proteins between the stationary and mid-exponential phases varied significantly between both conditions (p-value of < 0.05 and fold change of ± 1.25). For the aerobic cultures, 364 proteins were significantly more abundant under carbon starvation (stationary phase), while this only accounted for 78 proteins under anaerobiosis (**Figure 1c and Table 1**). A significantly lower abundance was also observed in the stationary phase for 174 proteins than in the exponential growth phase in the presence of oxygen and for 42 proteins only in the absence of oxygen (**Figure 1c and Table 1**).

**TABLE 1.** Number of proteins with significant changes during the transition to subsequent growth phase under aerobic and anaerobic conditions for the yeast control strain (CEN.PK113-7D). The total number of proteins indicates the number of proteins that were detected in at least two biological replicates. Proteins were normalized to the preceding growth phase. Only proteins with a fold change of 1.25 or greater (which corresponds to a log2 fold change of +/− 0.32) and a p-value of at least 0.05 are considered. The number of proteins quantified indicates the number of proteins that were detected and quantified in at least two biological replicates.

| CEN.PK113-7D growth transition | #Proteins quantified | # more abundant | # less abundant |
| --- | --- | --- | --- |
| **ANAEROBIC** |  |  |  |
| Late exponential / Mid-Exponential | 1092 | 1 | 22 |
| Early stationary / Late Exponential | 1092 | 52 | 3 |
| Stationary / Early Stationary | 1092 | 0 | 0 |
| Stationary / Mid-Exponential | 1092 | 78 | 42 |
| **AEROBIC** |  |  |  |
| Late exponential / Mid-Exponential | 998 | 12 | 21 |
| Early Diauxic / Late exponential | 998 | 67 | 4 |
| Mid-Diauxic / Early Diauxic | 1168 | 125 | 24 |
| Stationary / Mid-Diauxic | 1168 | 90 | 34 |
| Stationary / Mid exponential | 1168 | 364 | 174 |

Deprived of usable carbon source, stationary-phase yeast cells generally arrest growth, thereby entering a state of decreased metabolism and biosynthesis and yielding overall lower transcription and translation rates [43, 44]. Ribosomal proteins have been shown to be expressed at lower levels in the stationary phase [45, 46]. In good agreement with physiological data, the proteins involved in processes associated with protein synthesis and cellular growth showed decreased abundance in the transition between the exponential and stationary growth phases under both aerobic and anaerobic conditions, as shown in the categories ‘gene expression’, ‘ribosome assembly’ and ‘cellular macromolecule biosynthetic process’ (**SI Tables 6 and 7**). Yeast cells transition from respiro-fermentation on glucose to full respiration using ethanol as a primary carbon source in the presence of oxygen. This increase in respiratory activity was well reflected in the proteome in this study, as proteins more abundant in the stationary phase (than in the exponential phase) were typically associated with mitochondrial respiration in the aerobic conditions, including ‘generation of precursor metabolites and energy’, ‘mitochondrion organisation’ and ‘transmembrane transport’ (**SI Table 6**). As expected, this response was not observed in the nonrespiring, anaerobic cultures. In these cultures, most proteins involved in carbohydrate catabolic and disaccharide metabolic processes showed an increased abundance in the stationary compared to the mid-exponential phase, presumably to ensure survival in growth-arrested cells. Proteins in the cellular components involving categories such as ‘cell periphery’ and ‘plasma membrane’ were also found to be more abundant (**SI Table 7**).

The comparison of the proteomic data across the growth phases revealed that the diauxic shift had the strongest impact on proteomic rearrangement, with 24 proteins with lower abundance and 125 proteins with higher abundance between the beginning of the diauxic growth and mid-diauxic phases (**Table 1 and SI Table 6**). The diauxic shift was characterised by an increased abundance in proteins involved in aerobic respiration, fatty acid metabolism and precursor metabolite and energy generation, in line with the switch from respiro-fermentative to fully respiratory metabolism. Conversely, the set of proteins with decreased abundance during the diauxic shift was enriched for proteins involved in protein synthesis in the cytosol. This result was also consistent with the decreased growth rate and thereby the protein synthesis rate of yeast cells grown on ethanol media as compared with glucose [47]. Under anaerobiosis, most proteomic changes occurred in the transition between exponential and stationary growth (55 proteins; i.e. 46% of all detected changes in abundance throughout the phases). Notably, prolonged cultivation during the stationary phase under anaerobiosis did not further alter the proteome (**Table 1 and SI Table 7**).

### Impact of oxygen on the proteomic rearrangements in the central carbon metabolism across the growth phases

The central carbon metabolism (CCM) consists of key pathways required for the conversion of carbon sources into the 12 building blocks for the synthesis of cellular components and encompasses ca. 150 transport proteins and enzymes [25]. The flow of carbon and electrons via the CCM therefore responds to the carbon source nature and abundance. As oxygen availability dictates how much ATP molecules can be produced from the carbon source, the CCM also responds to oxygen availability. The proteins involved in the CCM are therefore expected to be considerably affected by glucose and oxygen availability. In our study, 101 out of 142 CCM proteins (**SI Table 8**) were successfully quantified in at least one of the main four conditions (**Figure 2a**). In the presence of oxygen, 71 proteins had significantly different abundance over the time-course, while in the absence of oxygen, only 18 significant changes were observed.

**FIGURE 2.**
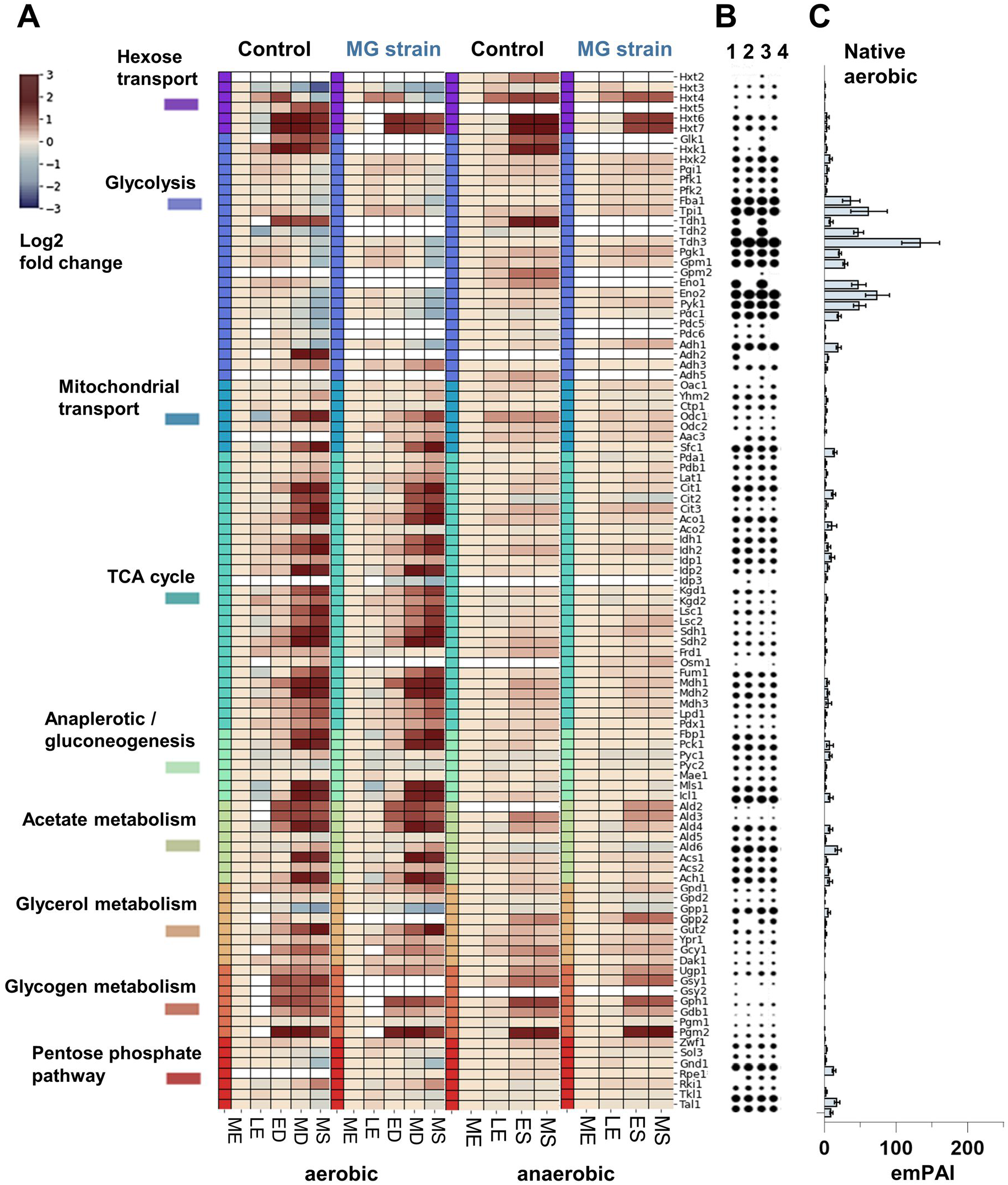
Yeast central carbon metabolism protein abundances under aerobic and anaerobic growth. A) The heat map shows the temporal log2 fold changes of the enzymes of the central carbon metabolism (CCM) of the control yeast CEN.PK113-7D and MG yeast for the mid-exponential (ME), late-exponential (LE), early-diauxic (ED), mid-diauxic (MD), early-stationary (ES), and mid-stationary (MS) phases, compared to the mid-exponential (ME) phase for each condition. The proteins belonging to specific pathways of the CCM are highlighted with different colours. White gaps in the map indicate that the protein was not detected, or that it has been deleted in case of the MG strain. No filtering for significance or fold-change thresholds was applied for this figure, and all enzymes that were detected were included in the heat map. B and C) The circle graph (1=control aerobic, 2=MG aerobic, 3=control anaerobic and 4=MG anaerobic) on the right express the emPAI values of the individual proteins for each condition as circle areas. The bar graph on the right shows the averaged emPAI values per enzyme. Standard deviations are indicated by the error bars in the bar graph.

*S. cerevisiae* harbours a set of 17 proteins able to transport hexoses, known as Hxt proteins. The expression of these proteins is primarily dictated by hexose (mostly glucose) abundance. Being membranebound and having low abundance, Hxt proteins are typically difficult to detect in proteomic studies; their high level of homology makes their identification challenging. Nevertheless, six Hxt proteins were quantified in the present dataset: Hxt2, 3, 4, 5, 6 and 7. Four of these Hxt proteins could be quantified in all samples, irrespective of strain and oxygen supply. Hxt6 and Hxt7 share a high protein sequence similarity (>99%) and were therefore considered as one protein group in this study. These were the most abundant Hxt proteins and were consistently more abundant upon glucose exhaustion in all tested conditions (**Figure 2a**), in good agreement with their high affinity for glucose [48]. Hxt3 and Hxt4, also identified in all conditions, responded differently in the presence and absence of oxygen. The abundance of Hxt4, a high-affinity transporter, expectedly increased upon glucose exhaustion but decreased upon reaching the stationary phase in the aerobic cultures; meanwhile, it remained high under anaerobiosis. The low-affinity transporter Hxt3 was detected at high and low glucose concentrations [49] but decreased in abundance across the growth curve and most significantly in the aerobic stationary phase. Hxt5 was only detected in the control strain in the presence of oxygen, but its abundance was in line with its induction by non-fermentable carbon sources and decreasing growth rates [48]. Hxt2 was only detected in the anaerobic cultures of the control strain. Thereby, Hxt2 increased in abundance upon glucose exhaustion, as expected for a high-affinity transporter.

Among the 26 glycolytic and fermentation enzymes, 13 major isoforms are constitutively expressed with high abundance, while the remaining are minor isoforms with lower abundance and condition-dependent expression [26, 50]. This notion was well reflected in the present comprehensive dataset, in which 23 of these proteins could be quantified in both conditions of the control strain. All major isoenzymes were found; their abundance remained constant under anaerobiosis but slightly decreased in the stationary phase under aerobiosis (**Figure 2a**). The majority of the minor isoenzymes were detected in at least one of the conditions. The minor glyceraldehyde-3P dehydrogenase Tdh1 was found to be expressed with and without oxygen. However, the enzyme was generally more abundant upon glucose exhaustion. Similarly, Glk1 and Hxk1, glucose-repressed isoenzymes of the predominant hexokinase 2, were also more abundant upon glucose exhaustion. Adh2, alcohol dehydrogenase repressed by glucose and induced by ethanol [51], was only detected in the presence of oxygen, and its abundance strongly increased in the mid-diauxic phase. Interestingly, the estimated total amount of the glycolytic enzymes was larger than the sum of the other CCM enzymes in all conditions (**Figure 2b and c**).

The TCA cycle, encompassing a set of 22 proteins located in the mitochondrion, is particularly active during respiration. The majority of the TCA cycle proteins were detected in our experiments. Their abundance generally strongly increased during the diauxic shift but did not change under the anaerobic conditions (**Figure 2a**). A small group of proteins remained unaffected during progression through the growth phase under anaerobiosis. For example, Pda1, Pdb1 and Lat1, which are three of the five subunits of the pyruvate dehydrogenase complex, Aco2 and Idp1 remained unchanged. Interestingly, Frd1 and Osm1, which are important for protein folding in anaerobic conditions [52], also remained unaffected. Many CCM metabolites cross the mitochondrial membrane via transporter proteins, and increased respiratory activity is expected to increase the flux of metabolites between the cytosol and mitochondria. Accordingly, the abundance of the seven quantified mitochondrial transporters increased after glucose exhaustion aerobically but not anaerobically, with a marked increase for Odc1 and Sfc1, which are carboxylic acid antiporters. Despite the increase in respiratory activity during the diauxic shift and phase, which was reflected in the TCA cycle proteins, the abundance of anaplerotic proteins Pyc1, Pyc2 and Mae1 remained unchanged. Conversely, the abundance of the gluconeogenic proteins Fbp1 and Pck1 and glyoxylate cycle proteins MLs1 and Icl1 required for ethanol utilisation strongly increased upon glucose exhaustion aerobically but was expectedly very stable anaerobically. Growth on non-fermentable carbon sources requires a complex metabolic rearrangement to supply cytosolic and mitochondrial acetyl-CoA. The expression of the proteins involved in acetate and acetyl-CoA metabolism, particularly Acs1 and Ach1, accordingly increased during the diauxic phase but was not visibly affected by glucose exhaustion under anaerobiosis.

Redox metabolism, a key for cell survival, is balanced according to oxygen availability. While respiring cells can oxidise the NADH produced during glucose assimilation via oxidative phosphorylation, two-step glycerol formation from dihydroxyacetone phosphate is the major electron sink in the absence of oxygen. The abundances of paralogues Gpd1, Gpd2, Gpp1 and Gpp2 were not strongly affected across the different phases and growth conditions (**SI Figure 3a**), which is in agreement with the reported transcriptional regulation and (in)activation by post-translational modification (phosphorylation) [53]. The only noticeable changes were the decreased abundance of Gpp1 in the presence of oxygen and increased abundance of Gpp2 upon glucose exhaustion in the absence of oxygen. In aerobic conditions, NADH is generally oxidised by external or internal NADH dehydrogenases, which shuttle the electrons into the mitochondrial electron transfer chain. Contrarily, anaerobic yeast cultures reoxidise the excess NADH formed during biosynthesis via glycerol production [54]. The increased abundance of Gut2 during the diauxic shift under aerobiosis can be explained by both its role in glycerol utilisation and redox balance maintenance (**SI Figure 3a**).

### Oxygen-dependent dynamics in other pathways

Respiration is an important mechanism for energy conservation in the presence of the electron acceptor oxygen. After the diauxic shift, respiration becomes the main ATP source for the cells. Accordingly, the abundance of the ATP synthases and cytochrome oxidases in the oxidative phosphorylation pathway increased significantly after glucose exhaustion (**SI Figure 6**). However, the abundance of these proteins remained constant over the entire growth curve in the anaerobic cultures. Respiring cells are prone to generation of reactive oxygen species (ROS), for instance by the production of superoxide during electron transfer during oxidative phosphorylation, which can induce the expression of stress tolerance genes [55]. Accordingly, Sod1 and Sod2, enzymes that detoxify superoxide and produce hydrogen peroxide [56], were significantly more abundant under the aerobic conditions than under the anaerobic conditions (**SI Figure 6**). Their abundance increased by at least twofold towards the stationary phase compared with that under log growth in the presence of oxygen, while the protein abundance remained constant under anaerobic growth. A similar protein profile was found for the peroxisomal and mitochondrial catalase A (Cta1) that detoxifies hydrogen peroxide. Nevertheless, the abundance of Ccs1, copper ion chaperone to Sod1, was relatively constant and similar between both conditions in the control strain (**SI Figure 6**).

Heme synthesis, encompassing a set of eight Hem proteins, also depends on oxygen availability [57]. In this study, seven Hem proteins could be quantified. The protein abundance of Hem1,2, 3, 12 and 15 was either similar between the aerobic and anaerobic conditions or higher aerobically (**Figure 3a**). However, Hem13 and Hem14 were only quantified in the anaerobic conditions, although Hem14 could be quantified only in one biological replicate based on a few peptides. Hem13 was confidently quantified with more than 10 peptides for each biological anaerobic replicate but was not found aerobically. Surprisingly, both Hem13 and Hem14 require oxygen for enzymatic activity.

**FIGURE 3.**
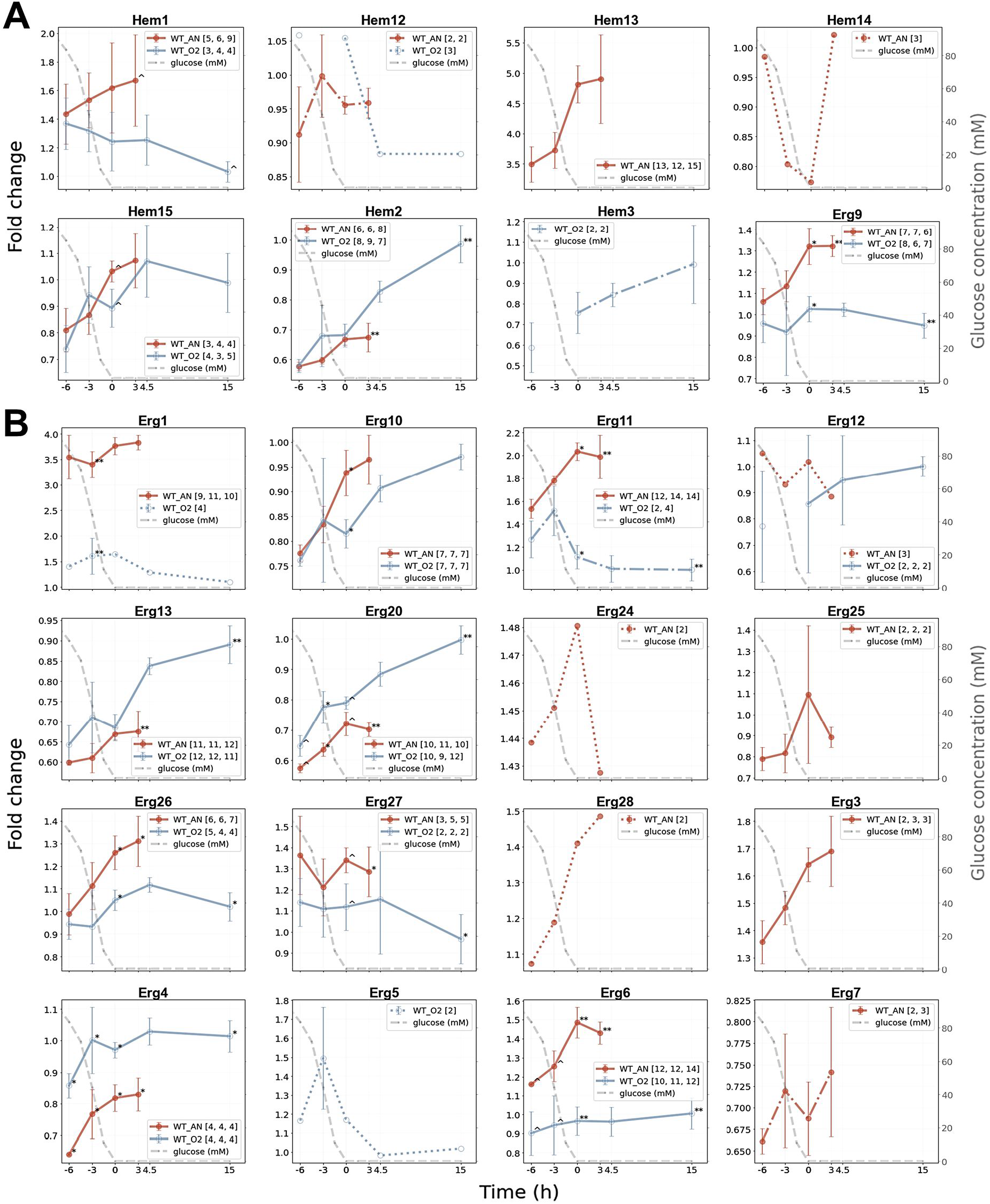
Protein fold change line graphs for non-respiratory oxygen-dependent pathways during aerobic and anaerobic growth. The foldchange values were plotted against the time relative to glucose depletion (t=0) in hours for proteins involved in heme (A) and sterol (B) synthesis. The different colours of the line graphs represent: “orange” the control yeast strain under anaerobic conditions (WT_AN), “light blue” the control strain under aerobic conditions (WT_O2). The line graphs represent the average of the three biological replicates, where the error bars indicate the standard deviation. The grey dashed line represents the glucose concentration over time (mM, secondary y-axis). The number of quantified peptides per biological replicate are shown in brackets. Asterisks (*) and circumflexes (^) indicate the significance (p values) between the aerobic and anaerobic experiments as follows: p< 0.001 (***), p< 0.01 (**), p< 0.05 (*), and p< 0.1 (^).

Finally, sterol synthesis also requires oxygen and ergosterol and is therefore supplied as an anaerobic growth factor during anaerobic yeast cultures [58]. Herein, 17 Erg proteins were found in either the aerobic or anaerobic condition or both (**Figure 3b**). The Erg proteins that need oxygen, such as Erg1, Erg3, Erg11 and Erg25–28, were either solely present anaerobically or more abundant in the anaerobic cultures than in the aerobic cultures.

### Survival in the stationary phase in response to oxygen availability

A previous study has shown a strong effect of oxygen availability and the presence of transition through the diauxic phase on yeast cell robustness during the stationary phase [59]. This work proposed that oxygen availability had a positive effect on the adenylate energy charge, longevity, stress response and thermotolerance during the stationary phase. However, this study was based on changes in the transcript levels, without confirmation at the protein level. In particular, limited data are known regarding the proteome dynamics under anaerobic conditions. To fill this knowledge gap, we studied the proteomic differences between the stationary phase in the anaerobic and aerobic conditions. A total of 249 proteins were significantly more abundant, and 125 were less abundant in the presence of oxygen than in the absence of oxygen (p-value of < 0.05 and fold change of ± 1.25; **SI Table 9 and SI Figure 4**). In the stationary phase, the aerobic cells still relied on respiration, and the proteins involved in respiration were accordingly more abundant under the aerobic conditions. Conversely, the yeast cells entered the stationary phase rather abruptly after glucose depletion under the anaerobic conditions. The proteins associated with ‘biosynthesis’, ‘glycolysis’ and ‘cytoplasmic translation’ were more abundant in the anaerobic stationary cells than in the aerobic cells. Several ribosomal proteins were also more abundant supposedly owing to the lack of time and resources required to adjust to the altering conditions. Furthermore, the proteins involved in storage metabolism, in particular glycogen metabolism, were enriched anaerobically.

Yeast cells generally accumulate storage carbohydrates in sugar-rich conditions that can be used as carbon and energy source to ensure survival during the stationary phase. Anaerobic cultures are entirely dependent on glycogen and trehalose as energy storage components. Conversely, the presence of oxygen enables yeast cells to metabolise other available nutrients, such as lipids and amino acids [60–62]. In this study, several proteins involved in glycogen metabolism were more abundant in the anaerobic cultures than in the aerobic cultures (**SI Figure 3b**). Glucose-6P is the starting point for glycogen metabolism and is converted into glucose-1P by Pgm1 or Pgm2. Both proteins were detected under the aerobic and anaerobic conditions, although Pgm2 was detected with higher coverage and confidence in each biological replicate, as it is the major isoform. The abundance of Pgm2 increased strongly (approximately 2.8-fold) after glucose depletion under the anaerobic conditions. Unfortunately, Pgm2 was only detected in the aerobic conditions after glucose exhaustion, although it was less abundant than that in the anaerobic conditions. Glucose-1P is subsequently converted to UDP-glucose by Ugp1. This enzyme was consistently more abundant in the absence of oxygen over the entire growth curve. Nevertheless, the protein profile was similar under both conditions, as the abundance of Ugp1 increased considerably after glucose was depleted. In the following step, glycogen synthesis is initiated by Glg1 and Glg2, which were not found in our analysis. Glycogen is subsequently generated by Gsy1/2, where only Gsy1 was detected in our experiments with sufficient confidence in the anaerobic cultures. Glycogen is a polymer that can be branched by Glc3; however, this enzyme was not quantified in our study. Glycogen can also be utilised by Gph1 to form glucose-1P again or by Gdb1 for conversion to glucose. Herein, only Gph1 showed a profile comparable to that of Pgm2.

Trehalose is another storage metabolite in yeast, which is synthesised from UDP-glucose. It is converted into trehalose-6P by Tps1 and subsequently into trehalose by Tps2. Trehalose is utilised by Nth1, Nth2 and Ath1 and converted into glucose again. In our study, the enzymes leading up to trehalose had similar protein profiles under the aerobic and anaerobic conditions, and the protein abundance increased significantly after glucose depletion (**SI Figure 3b**). However, utilisation of trehalose was more difficult to capture, as only Nth1 was quantified with only a few unique peptides under the aerobic and anaerobic conditions. Furthermore, in the absence of oxygen, yeast cells could not catabolise fatty acid by beta-oxidation as energy reserve during carbon starvation. Proteins such as Fox2, Pox1, Cat2 and Crc1 showed relatively constant expression levels over the anaerobic growth curve, while their abundance increased drastically after glucose depletion in the presence of oxygen (**SI Figure 6**).

Finally, aerobic stationary-phase cells are known to acquire increased robustness and stress tolerance during transition to the stationary phase. However, previous studies have indicated that this does not apply to anaerobic cultures to the same extent [59]. Stress proteins include a range of heat shock proteins (Hsp) with various functions. The fold changes and levels of Hsp were comparable in the presence and absence of oxygen in the exponential growth phase (**SI Figure 5**), but Hsp was more abundant in both aerobic and anaerobic conditions towards the end of the growth curve in the MS phase. This increase was markedly greater in the aerobically cultured cells, resulting in a significantly lower anaerobic fold change of the proteins in the stationary phase.

### Proteome-level alterations following genetic minimisation of the glycolytic pathway

The genetic reduction in the MG strain consisted of the removal of the 13 minor enzymes involved in glycolysis and fermentation, only leaving the 13 major isoenzymes. The present dataset showed that most minor isoenzymes were expressed and quantifiable in all samples from the aerobic and anaerobic batch cultivations, with the exception of Gpm2 and Adh5 detected only in the anaerobic cultures, Adh2 detected only in the aerobic cultures, and Adh4, Gpm3 and Pyk2 not detected at all (**Figure 4**). Tdh2, Eno1 and Adh2 were abundant in the batch cultures, and deletion of the minor isoenzymes might therefore affect the yeast physiology. The MG strain was previously well characterised by physiological and transcriptome analyses in the presence of oxygen, revealing that the physiology and transcriptome of this strain was nearly identical to that of the control strain with a full set of glycolytic and fermentation genes. However, the proteome of the MG strain was not explored, and limited data are known regarding the response of the MG strain to anaerobiosis. The fluxes through glycolysis and the fermentation pathway are substantially higher in anaerobiosis than in aerobiosis [63], and the deletion of all minor isoenzymes might therefore have a different cellular impact in the presence and absence of oxygen. As previously observed, the physiology of the MG and control strains in the aerobic and anaerobic cultures was nearly identical (**SI Table 1**) [26].

**FIGURE 4.**
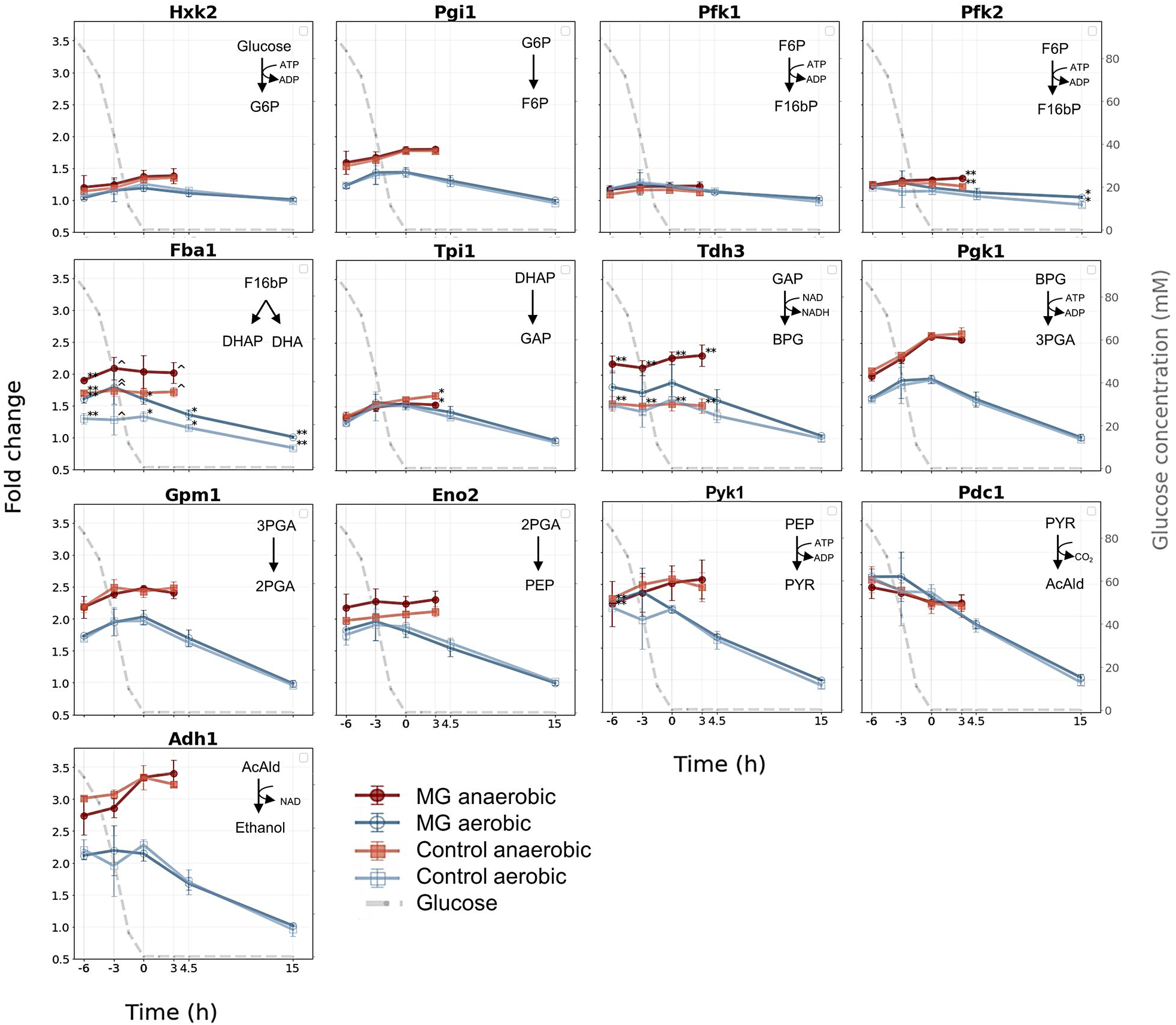
Protein fold change line graphs for the major glycolytic enzymes under aerobic and anaerobic growth. The fold-change values were plotted against the time relative to glucose depletion (t=0) in hours for all proteins of the major glycolytic enzymes. The different colours of the line graphs represent: “red” for the MG strain under anaerobic conditions, “dark blue” the MG strain under aerobic conditions, “orange” for the control yeast strain under anaerobic conditions and “light blue” for the control strain under aerobic conditions. Shown are the average values of the biological triplicates where the error bars indicate the standard deviations. The grey dashed line represents the glucose concentration over time (mM, secondary y-axis). Asterisks (*) and circumflexes (^) indicate the significant changes between the control and MG strain. P-values are indicated as follows: < 0.001 (***), < 0.01 (**), < 0.05 (*), and < 0.1 (^).

The difference in the abundance of the glycolytic and fermentation proteins between the MG and parental control strains was assessed by comparing the expression levels across the entire growth curve using a two-sided two-sample t-test. Remarkably, all major isoenzymes displayed identical time profiles between the MG and control strains; they were very stable under anaerobiosis, but their profiles decreased after the mid-diauxic phase under aerobiosis (**Figure 4**). With the exception of Fba1 and Tdh3, the abundance of the glycolytic and fermentation proteins was well conserved between the MG and control strains. The abundance of Tdh3 was consistently higher across all growth phases by 40–50% in the MG strain compared to the control strain anaerobically (p-value of < 0.01). The estimated protein amount of Tdh2 in the control strain was approximately one-third of the estimated protein amount of Tdh3 both under the aerobic and anaerobic conditions (**Figure 5**). The Tdh1 levels also markedly increased after glucose depletion in the control strain (**Figure 2a**). The loss of these relatively abundant minor isoenzymes and subsequent overall reduction of glyceraldehyde-3P activity in the MG strain might have caused a cross-regulation and an increased abundance of Tdh3. Interestingly, while the minor isoenzyme Eno1 was also abundant in the control strain (**Figure 5**), its deletion had no visible effect on the Eno2 level in the MG strain (**Figure 4**). The abundance of Fba1 was slightly (approximately 20–40%) but significantly higher in the MG strain than in the control strain both in the presence and absence of oxygen. Fba1 does not have isoenzymes, and this difference in abundance could therefore not be attributed to cross-regulation. Fba1 is an abundant protein in yeast, operating far from saturation [63]; the flux through glycolysis did not increase in the MG strain compared with that in the control strain. This increase in the abundance of Fba1 is therefore not likely explained by the need for a higher aldolase capacity in the MG strain.

**FIGURE 5.**
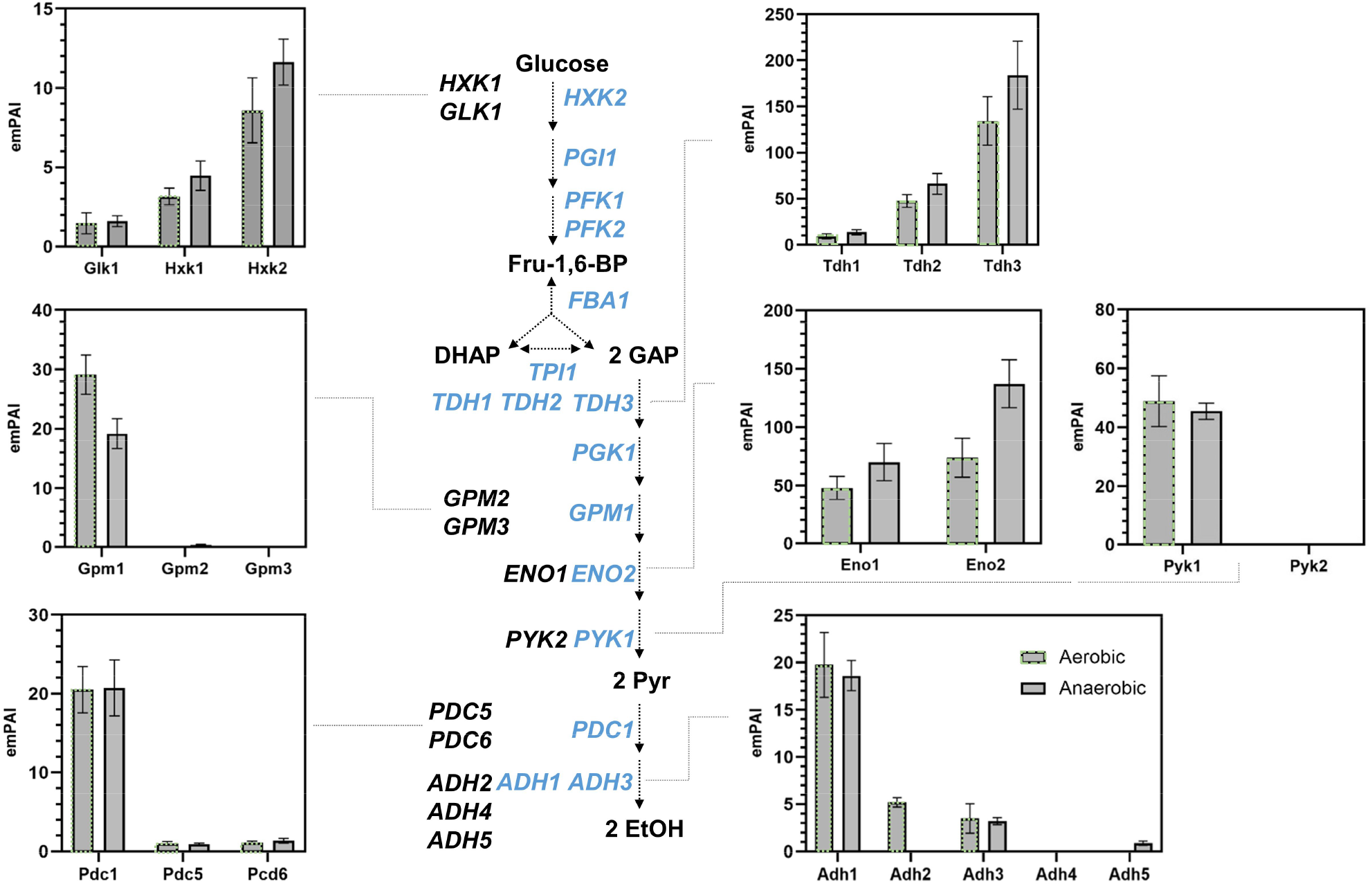
Abundance bar graphs for the glycolytic isoenzymes under aerobic and anaerobic conditions for the yeast control strain (CEN.PK113-7D). The bar graphs show the averaged protein abundances for the observed glycolytic isoenzymes expressed by their emPAI (exponentially modified protein abundance) indices, under aerobic (grey bars with green dashed lines) and anaerobic (grey bars) growth. The bars represent the average values of the individual biological replicates (with at least one identification per replicate), where the error bars indicate the standard deviation.

The enzyme level adjustments within the glycolytic and fermentation pathways following the deletion of the minor isoenzymes were remarkably small but slightly more pronounced under the anaerobic conditions. In line with this subtle phenotype, the proteome was not visibly affected by the deletion of the minor isoenzymes [cut-off p-value of 0.05 (5%), fold change threshold of ± 1.25; **Table 2 and SI Tables 10 and 11**]. Consequently, no enriched or depleted GO terms or KEGG pathways could be identified. The metabolic comparability between both strains is therefore underlined by their similar CCM enzyme (**Figure 2a**) and global proteome profiles.

**TABLE 2.** Number of proteins with significant changes between the yeast control strain (CEN.PK113-7D) and minimal glycolysis (MG) stain under aerobic and anaerobic conditions. The protein abundances of the MG strain were normalized to the yeast control strain (for the same growth phase and the same condition). Only proteins with a fold change of 1.25 (log2 fold change of +/− 0.32) or greater and a p-value of at least 0.05 are considered. The number of proteins quantified indicates the number of proteins that were detected and quantified in at least two biological replicates.

| MG mutant vs. control yeast | #Proteins quantified | # more abundant | # less abundant |
| --- | --- | --- | --- |
| **ANAEROBIC** |  |  |  |
| Mid exponential | 978 | 10 | 13 |
| Late exponential | 978 | 10 | 8 |
| Early stationary | 978 | 8 | 18 |
| Stationary | 978 | 7 | 6 |
| **AEROBIC** |  |  |  |
| Mid-Exponential | 1019 | 18 | 10 |
| Late-Exponential | 950 | 33 | 13 |
| Early-Diauxic | 1019 | 15 | 9 |
| Mid-Diauxic | 1019 | 3 | 3 |
| Stationary | 1019 | 3 | 2 |

## DISCUSSION

To our knowledge, this study provides the most comprehensive proteomic study on the response of *S. cerevisiae* CEN.PK113-7D to oxygen and nutrient availability. We employed tightly controlled batch bioreactor cultures and performed biological triplicates combined with standardised sample preparation protocols to ensure high reproducibility and accuracy. Albeit different yeast proteome dynamics studies have been performed over the past decades [16, 18, 20, 64], no study has yet captured the complete spectrum of conditions using the same strain and the same highly controlled experimental setup. Moreover, the transition from the exponential to the stationary phase under anaerobic conditions has not yet been investigated. This study moreover quantified the impact of genetic minimisation of the glycolytic pathway on proteome resource allocation using the recently established MG strain [23]. The established dataset therefore covers the quantitative analysis of 54 individual proteomes, where approximately 40% of all yeast proteins and approximately 99% of the complete protein biomass have been captured. The high reproducibility was also highlighted by the strong similarities between the proteomes of the MG and control strains. This comprehensive and accurate dataset therefore provides an ideal resource for applied and fundamental studies in yeast and more particularly for *in silico* proteome allocation studies.

The most remarkable observation was the substantially smaller proteome response of yeast cells grown under anaerobiosis than under aerobiosis. In both conditions, yeast cells had to tune their metabolism to the transition from glucose excess to exhaustion, leading to a shift from exponential to stationary growth. However, these drastic changes triggered a far milder response than did the aerobic transition to and out of the diauxic phase, which represented 58% of the measured changes in protein abundance. During the diauxic shift, cells rewire the proteome for respiratory growth on ethanol as a main carbon source. This transition leads to a multitude of physiological and morphological changes, including smaller cell size, increased mitochondrial volume, decreased growth rate, increased respiration rate and therefore increased ROS production, induction of gluconeogenesis and the glyoxylate cycle, and large changes in the fluxes in CCM. These changes were well reflected by the observed changes in the proteome allocation in this study. For instance, an increase in mitochondrial protein abundance was observed, while the glycolytic proteins were simultaneously downregulated [13–15, 65]. The transition from the diauxic to the stationary phase led to a further strong modification of the abundance of approximately 120 proteins. Conversely, as few as 55 proteins showed strongly altered abundances after glucose depletion under anaerobiosis, and prolonged cultivation in the stationary phase did not further alter the proteome. Using the exact same strain and experimental setup, Bisschops *et al*. (2015) [66] showed a stronger and faster decrease in viability upon glucose exhaustion for anaerobic cultures than for aerobic cultures. Based on physiological and transcriptome data, the authors attributed this lack of robustness to the inability of cells to adapt to glucose exhaustion in the absence of oxygen. Conversely, the diauxic shift provides the time and resources needed to transition from fast growth to growth arrest in the presence of oxygen. The present proteomics study supports this view in different ways. The small protein response during transition from sugar excess to depletion suggests that the anaerobic cells do not have the means or proper regulatory network to adjust to the new conditions. Furthermore, aerobic cultures acquire robustness and stress tolerance during transition to the stationary phase as a result of the expression of the ‘stress-response’ genes [67], such as Hsp. Both aerobic and anaerobic cultures showed a similar ‘stress signalling’ as shown by the increase in the Hsp level towards the stationary phase; however, this increase was far less pronounced in the anaerobic cultures. The abundance of Hsp was therefore substantially lower in the absence of oxygen, in line with the lower transcription of Hsp and lower thermo-tolerance of the anaerobic stationary-phase cultures than of the aerobic cultures observed by Bisschops *et al*. [66]. Considering that industrial-scale processes favour anaerobic environments for practical and financial reasons, the present results provide valuable information for the construction of predictive metabolic models [5, 9, 68–73].

In the presence of oxygen, yeast cells switch from respiro-fermentative to full respiratory metabolism once glucose is depleted. Accordingly, the abundance of respiration-related proteins increased upon glucose exhaustion in the aerobic cultures herein, while their protein profiles remained constant in the anaerobic cultures. Several other non-respiratory pathways in *S. cerevisiae*, such as fatty acid beta-oxidation and haeme and sterol synthesis, are oxygen-dependent. Expectedly, most detected proteins in these pathways were aerobically more abundant or contained similar abundance profiles to the anaerobic cultures. Nevertheless, oxygen-dependent protein Hem13 involved in haeme synthesis was only confidently quantified under anaerobic conditions, and the lack of detection in the aerobic conditions suggests that Hem13 is far less abundant. The transcription of Hem13 is repressed by oxygen and haeme itself [74, 75]; therefore, this protein lacks repression in the absence of oxygen, and its abundance is thereby increased. Similar protein profiles under anaerobic conditions were previously found for Hem1, Hem14 and Hem15 [18]. Several oxygen-dependent Erg proteins were also more abundant or solely detected under anaerobiosis. Herein, the transcription of various Erg proteins was regulated by oxygen, and an absence increased the expression of these Erg proteins [76] [77].

Glycolysis and alcoholic fermentation are well-studied pathways that play an important role in sugar conversion in the industry. Earlier studies have shown that major glycolytic isoenzymes are abundant proteins whose expression remains relatively stable irrespective of the growth environment, although these measurements often rely on gene transcription data or enzyme assays that cannot distinguish between isoenzymes [26]. The present proteome dataset confirmed that the abundance of most major glycolytic isoenzymes decreased during the diauxic shift and further decreased during the stationary phase in the presence of oxygen. Conversely, their abundance was unaffected during transition to the stationary phase under the anaerobic environments. Aerobic cultures in the stationary phase therefore display substantially lower glycolytic enzyme levels than do anaerobic cultures. For instance, a 2.5- to 3.5-fold lower abundance was observed for Pgk1, Gpm1, Pdc1 and Adh1. While this difference in abundance is not expected to affect survival in the stationary phase, in which the glycolytic flux is extremely low or absent, it will influence the ability of stationary-phase cells to reach fast growth when transitioned to a sugar-rich medium. When exposed to anaerobic sugar excess, cells grown aerobically to the stationary phase have to allocate resources to increase the abundance of glycolytic enzymes and reach fast growth, while cells pre-cultured anaerobically do not. As glycolytic enzymes are, next to ribosomal proteins, the most abundant proteins, this aspect should be considered during the startup phase of anaerobic industrial fermentations and their modelling. The expression of the minor glycolytic isoenzymes is condition-dependent, and several of these isoenzymes are reported to have distinct functions, especially during changes in carbon source availability. The present dataset showed that the presence of oxygen only visibly affected the abundance of Eno1, Tdh1 and Hxk1, as their abundance was significantly higher after glucose depletion under anaerobic conditions than under aerobic conditions (**Figure 2**). While some minor isoenzymes had a substantial abundance in yeast (**Figure 5**, e.g. Eno1, Tdh2 and Hxk1), with the exception of Tdh3, their removal did not trigger visible changes in the abundance of the major isoenzymes. Tdh3, glyceraldehyde dehydrogenase major isoenzyme, notably increased by 1.5-fold in the MG strain as compared to the control strain under anaerobic conditions (**Figure 4**). The same trend was observed aerobically, albeit much less pronounced. Similar to most glycolytic enzymes, glyceraldehyde dehydrogenase operates at overcapacity, meaning that the enzyme capacity largely exceeds the flux catalysed *in vivo* [78]. Therefore, the increased abundance of Tdh3 in the MG strain does most likely not result from the need to compensate for Tdh1 and Tdh2 deletion to maintain the glycolytic flux. Glyceraldehyde dehydrogenase isoenzymes do not have any well-described moonlighting functions. However, next to their cytosolic localization, they are also found in the cell wall in which they might play a yet uncovered role. The composition and structure of *S. cerevisiae* cell wall are affected by oxygen, and several cell wall proteins are specifically enriched under anaerobiosis (e.g. cell wall mannoprotein of the Srp1p/Tip1p family), which might explain the observed cross-regulation in the MG strain. Herein, Fba1 was also mildly but significantly upregulated in the MG mutant both aerobically (1.2 to 1.4-fold change) and anaerobically (1.15-fold change). As Fba1 does not have isoenzymes and is solely responsible for the glycolytic flux, its change in abundance is difficult to explain. Fba1 is also involved in vacuolar function as a subunit of the vacuolar V-ATPase [79]. However, as no or minimal differences were observed in the other components of V-ATPase between the MG and control strains, the molecular mechanism leading to the slightly higher abundance of Fba1 in the MG strain remains unclear. Many factors can alter the functionality of proteins, including post-translational modifications, protein localization or interactions with other proteins or biomolecules [80, 81]. A recent study has suggested that phosphorylation regulates the activity of many glycolytic enzymes [7]. However, the stable abundance of glycolytic proteins between the MG and control strains was well reflected in the stability of *in vitro* enzyme activity [26], suggesting the lack of differences in the post-transcriptional regulation between these strains. Taken together, remarkably few proteome-level changes were observed as a consequence of the genetic reduction of glycolysis. This similarity between both strains finally underscores the usefulness of the simplified MG strain for proteome allocation studies and for studying the role of post-translational modifications in the regulation of glycolysis.

The complete proteome dynamics and abundance data for the batch reactor-cultured CEN.PK113-7D strain and the related MG mutant for aerobic and anaerobic growth are shown in **SI Table 12**. Raw mass spectrometric data and unprocessed search files are publicly available via the PRIDE repository under the project code PXD031412.

## Supporting information

SI_word_doc

SI_overview_table

## AUTHOR CONTRIBUTIONS

MP, PDL and MDR designed the experiments. MDR performed experiments. MA helped with data generation. WB developed the proteome analysing pipeline in Python. MDR, WB and MP analysed the data. MDR, PDL and MP wrote the manuscript. All authors have given approval to the final version of the manuscript.

## NOTES

This work was supported by a TU Delft start-up fund. The Authors declare that there is no conflict of interest.

## ACKNOWLEDGMENTS

The authors are grateful to valuable discussions with our colleagues from the department of Biotechnology and acknowledge Carol de Ram, Christiaan Mooiman, Erik de Hulster, Jelle van Alphen and Casper van der Luijt for technical support.

