## Supplementary material for "Proteome dynamics during transition from exponential to stationary phase under aerobic and anaerobic conditions in yeast": SI_word_doc

### TABLE OF CONTENTS

SI Table 1: Biomass-specific substrate consumption and product formation rates — Page 2
SI Figure 1: Estimated absolute abundance of detected proteins — Page 3
SI Figure 2: Verification of reproducibility of biological and technical replicates — Page 4
SI Figure 3: Profiles of proteins involved in glycerol, glycogen and trehalose metabolism — Page 5
SI Figure 4: Volcano plots global proteome changes between aerobic and anaerobic conditions — Page 6
SI Figure 5: Profiles of proteins involved in glycerol, glycogen and trehalose metabolism — Page 7
SI Figure 6: Protein profile changes of selected pathways — Page 8

**SI Table 1: Biomass-specific substrate consumption and product formation rates of MG (IMX372) and control (CEN.PK113-7D) strain during aerobic and anaerobic exponential growth.** The average and standard deviations were calculated for three biological replicates per strain. The biomass specific consumption or formation rates were calculated for the substrate glucose ($q_s$), ethanol ($q_{EtOH}$), glycerol($q_{Glyc}$), acetate ($q_{Ace}$), carbon dioxide ($q_{CO2}$), and oxygen ($q_{O2}$). Significant differences ($p<0.01$) between control and MG yeast were calculated with an two-sided two-sample unpaired t-test and are highlighted with an asterisk. #The $qCO_2$ of IMX372 under anaerobic conditions is likely underestimated due to technical complications in the $CO_2$ off gas analysis equipment.

| | AEROBIC | | | | ANAEROBIC | | | |
|---|---|---|---|---|---|---|---|---|
| | CEN.PK113-7D | | IMX372 | | CEN.PK113-7D | | IMX372 | |
| | Average | Stdev | Average | Stdev | Average | Stdev | Average | Stdev |
| **$\mu^{max}$** ($h^{-1}$) | 0,413 | 0,012 | 0,396 | 0,002 | 0,366 | 0,003 | 0,355 | 0,012 |
| **$q_s$** ($mmol \cdot gDW^{-1} \cdot h^{-1}$) | -18,890 | 1,029 | -18,585 | 0,580 | -23,264 | 1,792 | -20,677 | 0,659 |
| **$q_{EtOH}$** ($mmol \cdot gDW^{-1} \cdot h^{-1}$) | 30,166 | 1,999 | 27,063 | 0,446 | 31,814 | 3,049 | 30,999 | 1,677 |
| **$q_{Glyc}$** ($mmol \cdot gDW^{-1} \cdot h^{-1}$) | 1,736 | 0,074 | 1,518 | 0,147 | 4,607 | 0,395 | 4,297 | 0,198 |
| **$q_{Ace}$** ($mmol \cdot gDW^{-1} \cdot h^{-1}$) | 0,494 | 0,082 | 0,493 | 0,080 | 0,560 | 0,092 | 0,540 | 0,045 |
| **$q_{CO2}$** ($mmol \cdot gDW^{-1} \cdot h^{-1}$) | 33,024 | 2,056 | 28,175 | 0,912 | 32,037 | 1,213 | 27,053*# | 1,343 |
| **$q_{O2}$** ($mmol \cdot gDW^{-1} \cdot h^{-1}$) | 9,479 | 0,533 | 7,978 | 0,149 | | | | |
| **RQ** | 3,517 | 0,173 | 3,574 | 0,215 | | | | |
| **$Y^{biomass/glucose}$** ($gDW \cdot gglucose^{-1}$) | -0,022 | 0,001 | -0,021 | 0,001 | -0,016 | 0,001 | -0,017 | 0,000 |
| **$Y^{ethanol/glucose}$** ($mol \cdot mol^{-1}$) | 1,598 | 0,111 | 1,458 | 0,025 | 1,379 | 0,032 | 1,503 | 0,129 |
| **$Y^{glycerol/glucose}$** ($mol \cdot mol^{-1}$) | 0,092 | 0,009 | 0,082 | 0,009 | 0,199 | 0,027 | 0,208 | 0,004 |
| **$Y^{acetate/glucose}$** ($mol \cdot mol^{-1}$) | 0,026 | 0,003 | 0,026 | 0,004 | 0,024 | 0,004 | 0,026 | 0,001 |
| **Carbon recovery** | 1,000 | 0,000 | 1,000 | 0,000 | 1,000 | 0,000 | 1,000 | 0,000 |

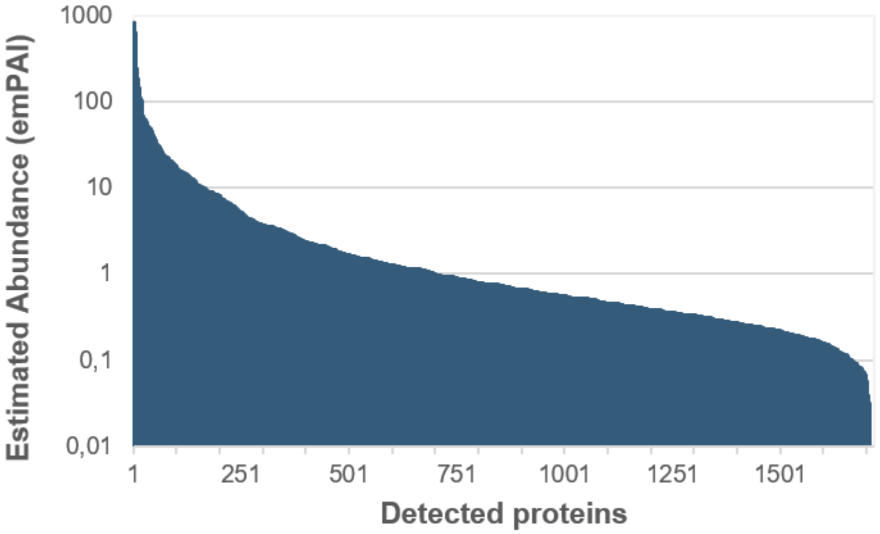


**SI Figure 1. Estimated absolute abundance of detected proteins across all experiments.** The absolute abundance was estimate using the Exponentially Modified Protein Abundance Index (emPAI) for all identified proteins across all experiments. The absolute protein amount is estimated by the number of sequenced peptides per protein. The abundance of each protein is an average of all experiments and detected in at least one biological replicate.

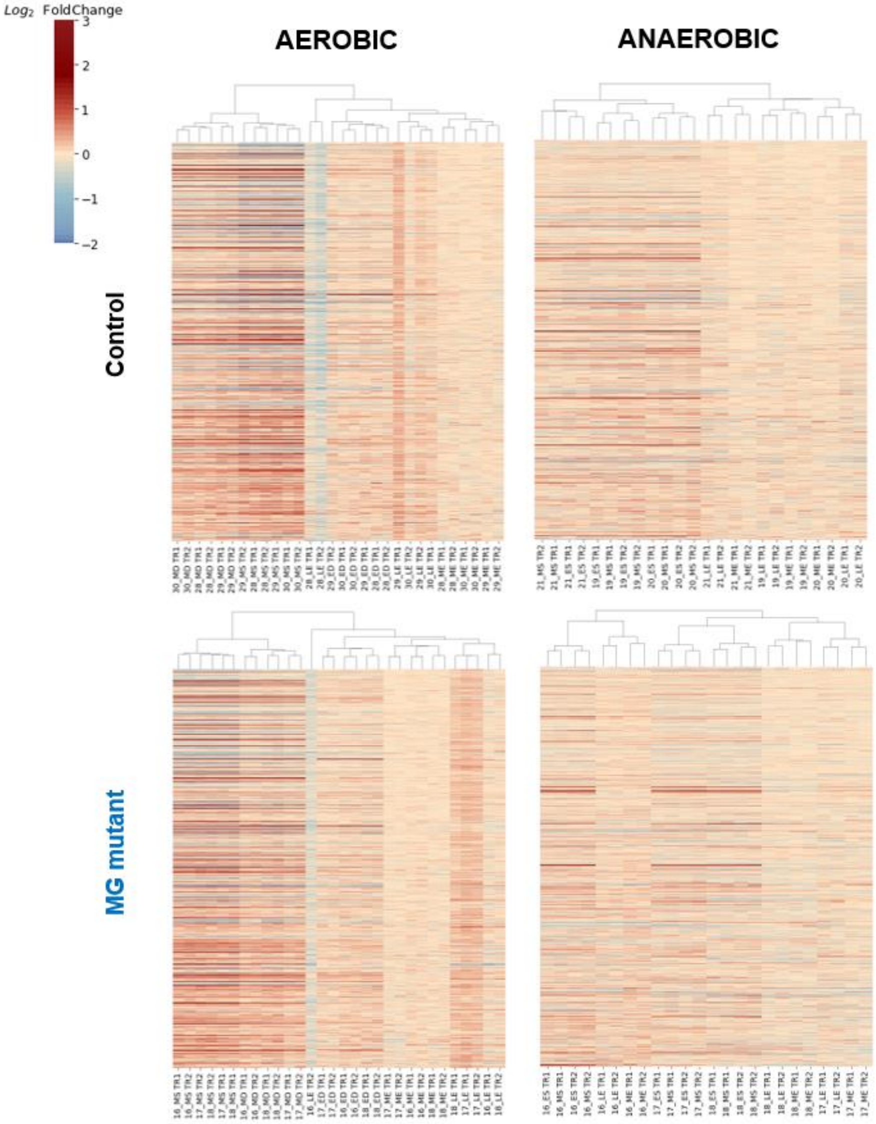


**SI Figure 2. Cluster analysis of abundance changes of proteins from the control and MG yeast strain under aerobic and anaerobic conditions.** The replicates were clustered based on Euclidean distances and with the average linkage method. The protein fold changes were normalised to the average of all the ME-phase experiments. The log2 of the fold changes was calculated and plotted so that no fold change (=0) is coloured white, a negative fold change is coloured blue and a positive fold change red. The columns labels represent the reactor number - indicating the biological replicate - the growth phase, and the technical replicate (TR) number. ME: mid-exponential, LE: late-exponential, ED: early-diauxic, MD: mid-diauxic shift, ES: early-stationary, MS: mid-stationary.

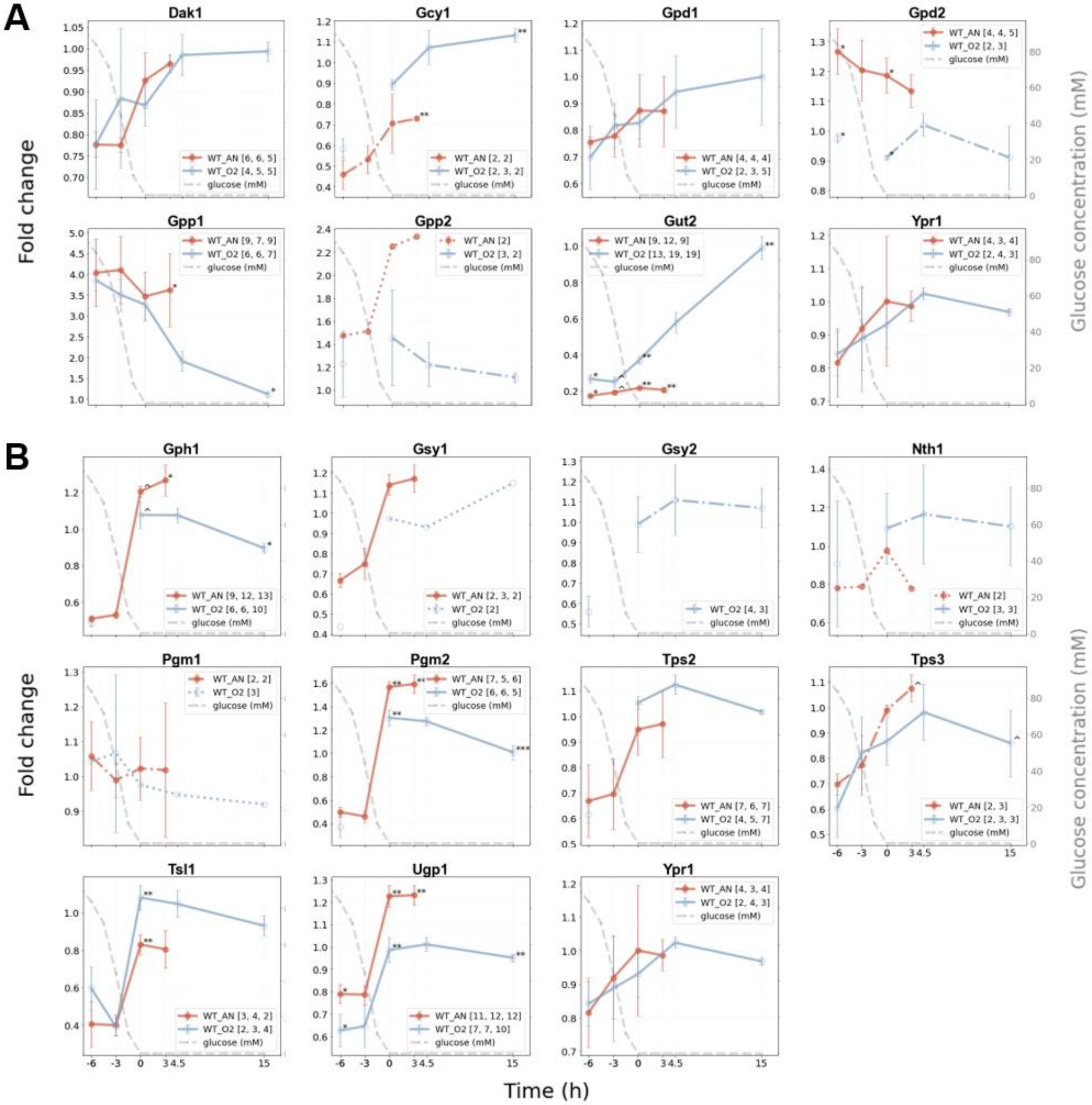


**SI Figure 3. Protein fold change line graphs for the glycerol, glycogen and trehalose metabolism during aerobic and anaerobic growth for the control yeast CEN.PK113-7D.** The biological-replicate-averaged fold-change values were plotted against the time relative to glucose depletion in hours for proteins involved in glycerol (A) and glycogen and trehalose (B) metabolism. The different colours of the line graphs represent: "orange" the control strain under anaerobic conditions (WT_AN) and "light blue" the control strain under aerobic conditions (WT_O2). The error bars show the standard deviation of the mean of the three biological replicates. The grey dashed line represents the glucose concentration over time (mM, secondary y-axis). The number of quantified peptides per biological replicate are indicated in brackets. Asterisks (*) and circumflexes (ˆ) indicate the significance between the aerobic and anaerobic experiments, which are as follows: p< 0.001 (***), p< 0.01 (**), p< 0.05 (*), and p< 0.1 (ˆ).

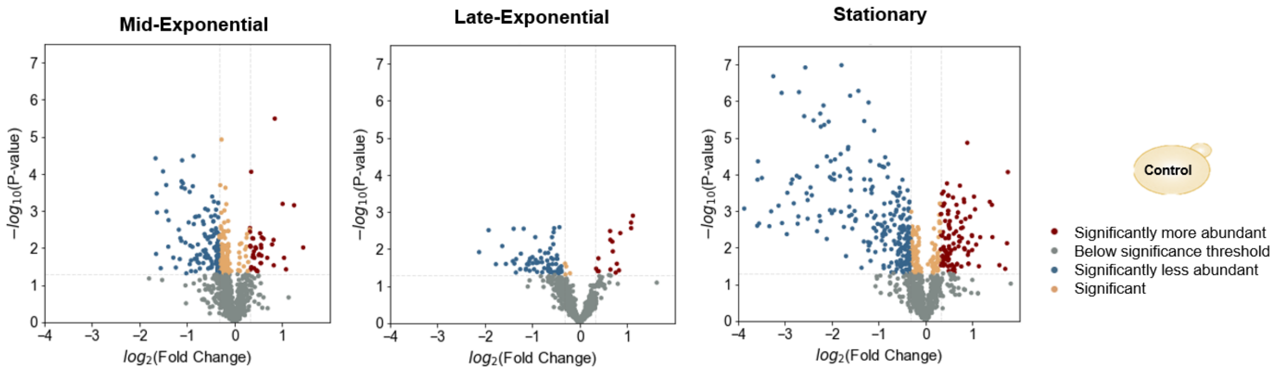


**SI Figure 4. Volcano plots showing the global proteome changes between aerobically and anaerobically cultured control yeast CEN.PK113-7D**. The log2 of the abundance fold change between the two conditions (normalised to the aerobic experiments) was plotted against the -log10 of the p-value. The mid-exponential (ME), late-exponential (LE) and mid-stationary (MS) phases were compared. P-value threshold <0.05, fold change threshold >1.5 (log2 fold change threshold +/- 0.32).

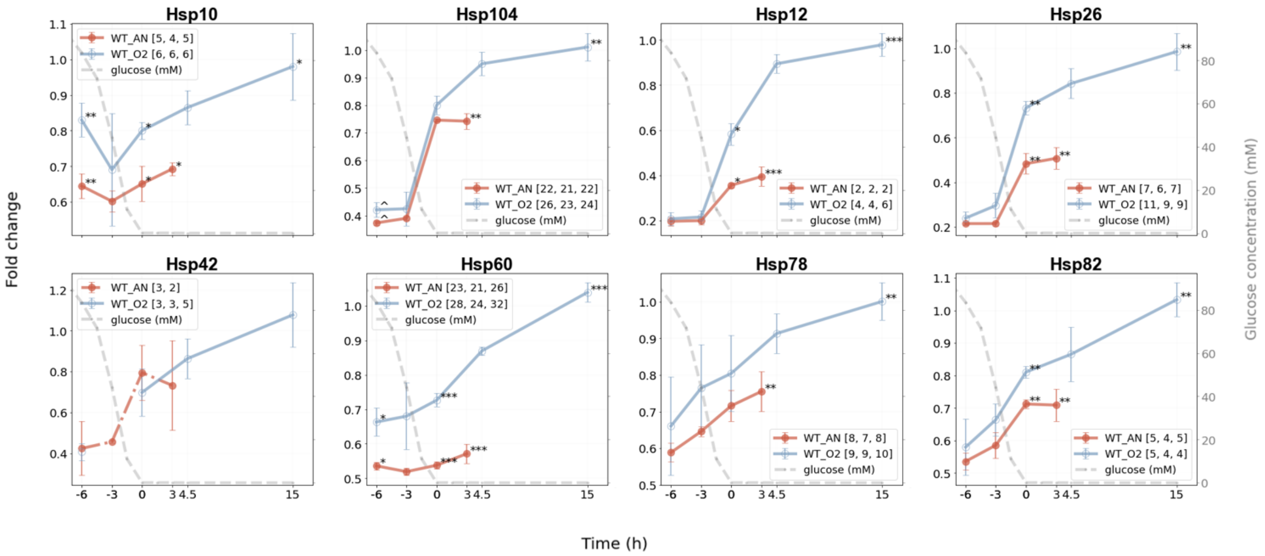


**SI Figure 5. Fold changes line graphs for heat shock proteins observed for the control yeast CEN.PK113-7D under aerobic and anaerobic conditions.** The biological-replicate-averaged FCs were plotted against the time relative to glucose depletion (0 h) in hours. Orange: CEN.PK113-7D under anaerobic conditions, light blue CEN.PK113-7D under aerobic conditions. The error bars show the standard deviation of the mean of the three biological replicates. In the legend, the numbers between brackets represent the number of unique peptides that were found in each biological replicate. The grey dashed line represents the glucose concentration over time (mM, secondary y-axis). Asterisks (*) and circumflexes (ˆ) indicate the significance between the aerobic and anaerobic experiments, which are: $p < 0.001$ (***), $p < 0.01$ (**), $p < 0.05$ (*), and $p < 0.1$ (ˆ).

### ATP synthesis

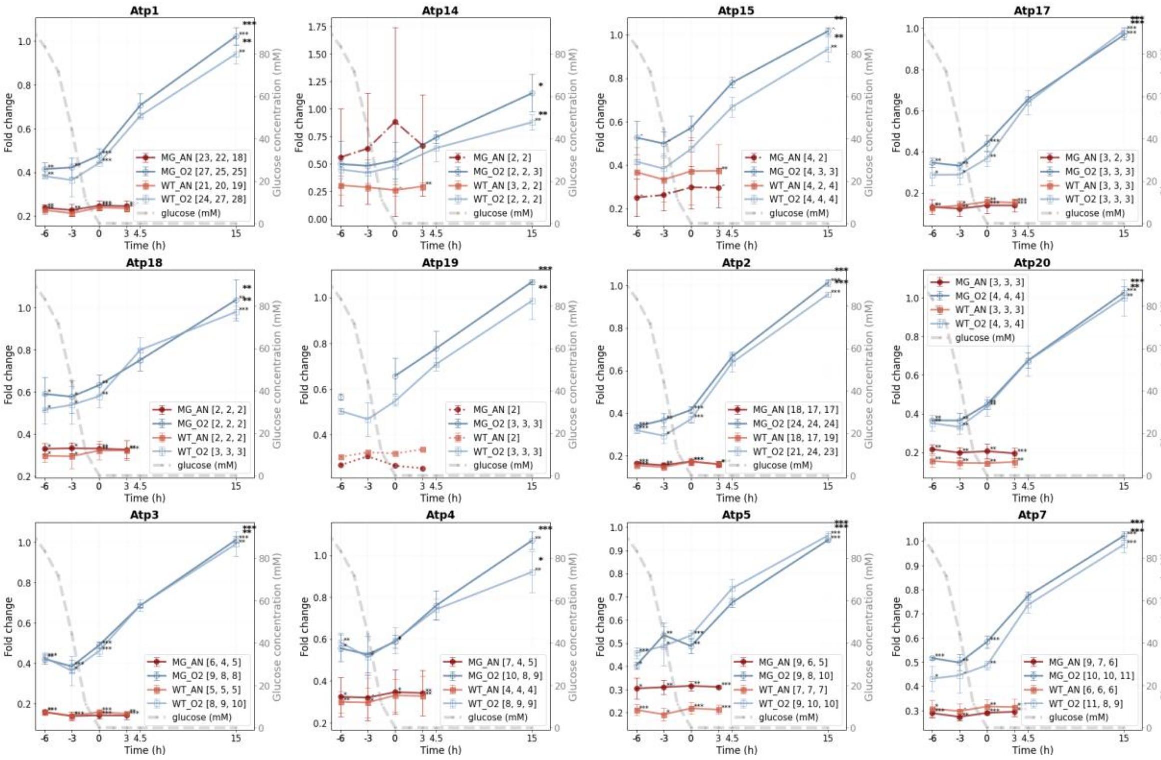


**SI Figure 6.1. Fold change line graphs for proteins of selected pathways for the control yeast and the MG strain under aerobic and anaerobic conditions.** The biological-replicate-averaged FCs of CEN.PK113-7D and IMX372 (MG) were plotted against the time relative to glucose depletion (0 h) in hours. Red: the MG strain under anaerobic conditions, dark blue: the MG strain under aerobic conditions, orange: the control strain under anaerobic conditions, light blue: the control strain under aerobic conditions. The error bars show the standard deviation of the mean of the three biological replicates. In the legend, the numbers between brackets represent the number of unique peptides that were found in each biological replicate. The grey dashed line represents the glucose concentration over time (mM, secondary y-axis). Asterisks (*) and circumflexes (ˆ) indicate the significance between either the (an)aerobic experiments (black annotation) or between the ME and MS phase. P-value levels are: $p < 0.001$ (***), $p < 0.01$ (**), $p < 0.05$ (*), and $p < 0.1$ (ˆ).

### Respiration

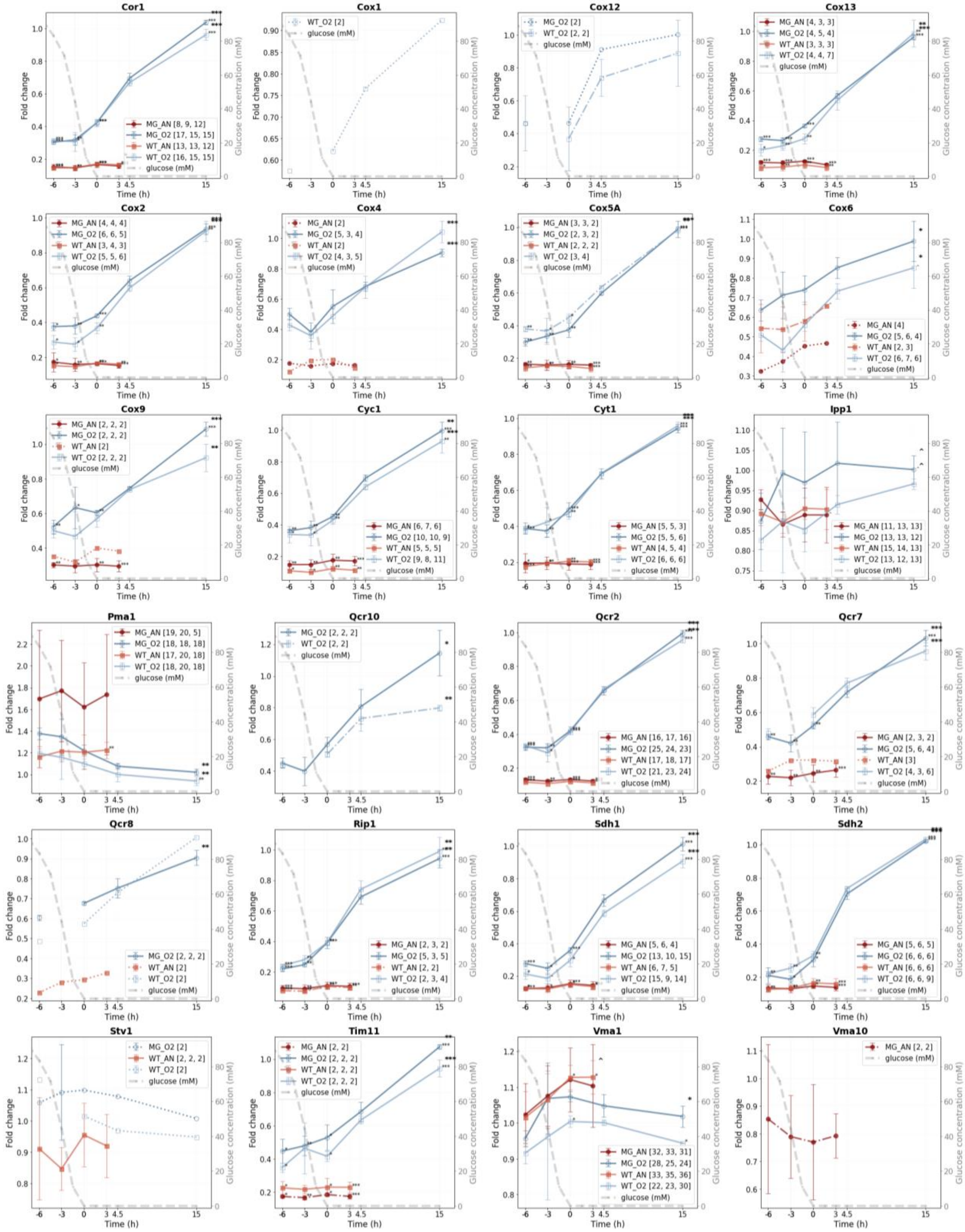

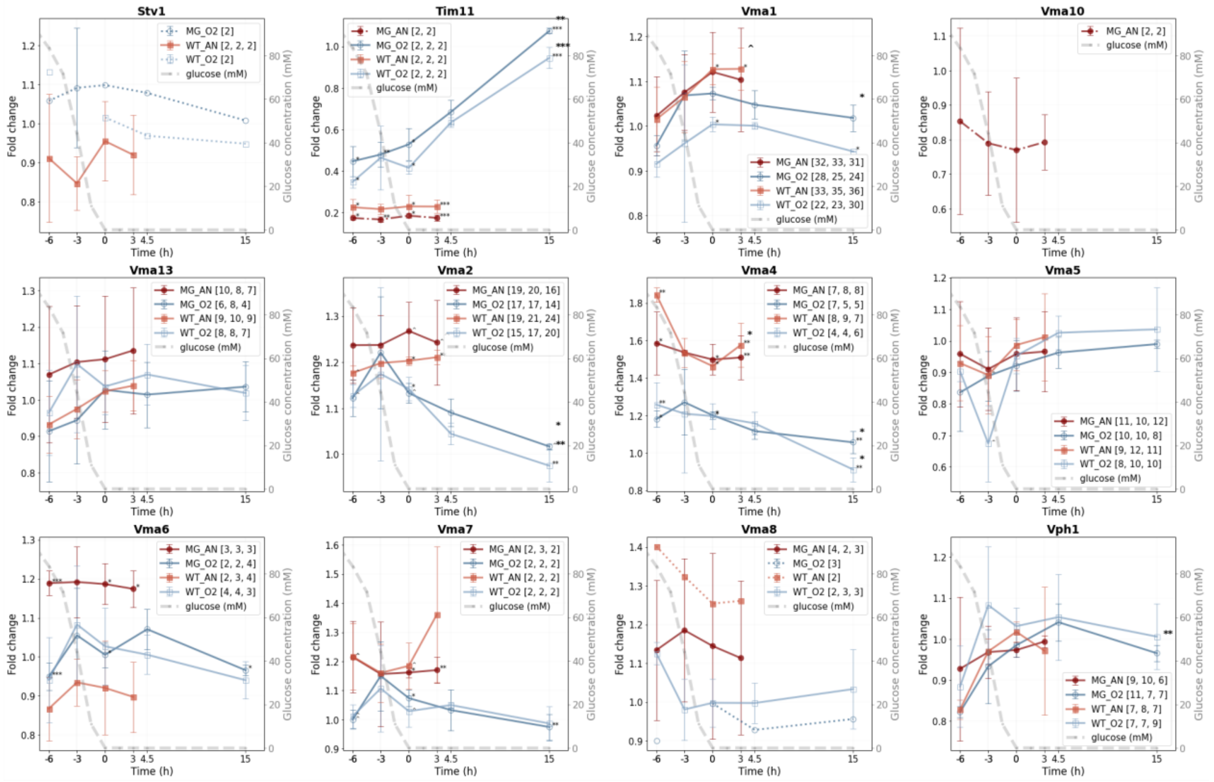


**SI Figure 6.2. Fold change line graphs for proteins of selected pathways for the control yeast and the MG strain under aerobic and anaerobic conditions.** The biological-replicate-averaged FCs of CEN.PK113-7D and IMX372 (MG) were plotted against the time relative to glucose depletion (0 h) in hours. Red: the MG strain under anaerobic conditions, dark blue: the MG strain under aerobic conditions, orange: the control strain under anaerobic conditions, light blue: the control strain under aerobic conditions. The error bars show the standard deviation of the mean of the three biological replicates. In the legend, the numbers between brackets represent the number of unique peptides that were found in each biological replicate. The grey dashed line represents the glucose concentration over time (mM, secondary y-axis). Asterisks (*) and circumflexes (^) indicate the significance between either the (an)aerobic experiments (black annotation) or between the ME and MS phase. P-value levels are: $p < 0.001$ (***), $p < 0.01$ (**), $p < 0.05$ (*), and $p < 0.1$ (^).

### Plasma membrane transport

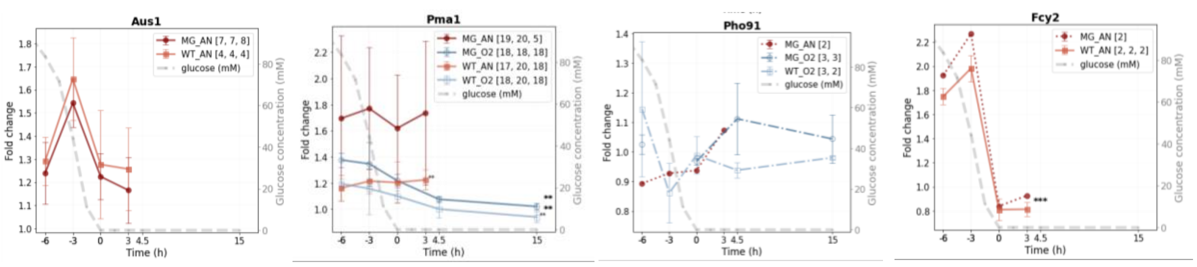


**SI Figure 6.3. Fold change line graphs for proteins of selected pathways for the control yeast and the MG strain under aerobic and anaerobic conditions.** The biological-replicate-averaged FCs of CEN.PK113-7D and IMX372 (MG) were plotted against the time relative to glucose depletion (0 h) in hours. Red: the MG strain under anaerobic conditions, dark blue: the MG strain under aerobic conditions, orange: the control strain under anaerobic conditions, light blue: the control strain under aerobic conditions. The error bars show the standard deviation of the mean of the three biological replicates. In the legend, the numbers between brackets represent the number of unique peptides that were found in each biological replicate. The grey dashed line represents the glucose concentration over time (mM, secondary y-axis). Asterisks (*) and circumflexes (^) indicate the significance between either the (an)aerobic experiments (black annotation) or between the ME and MS phase. P-value levels are: $p < 0.001$ (***), $p < 0.01$ (**), $p < 0.05$ (*), and $p < 0.1$ (^).

### Fatty acid metabolism

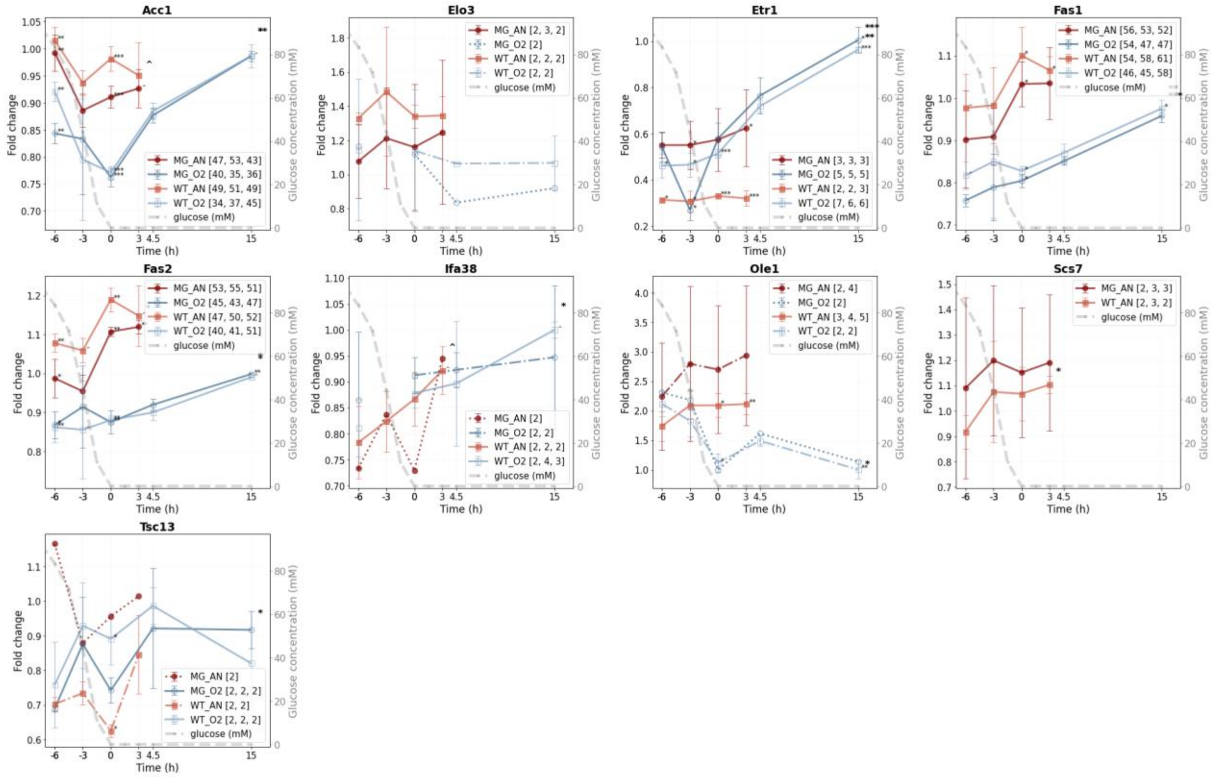


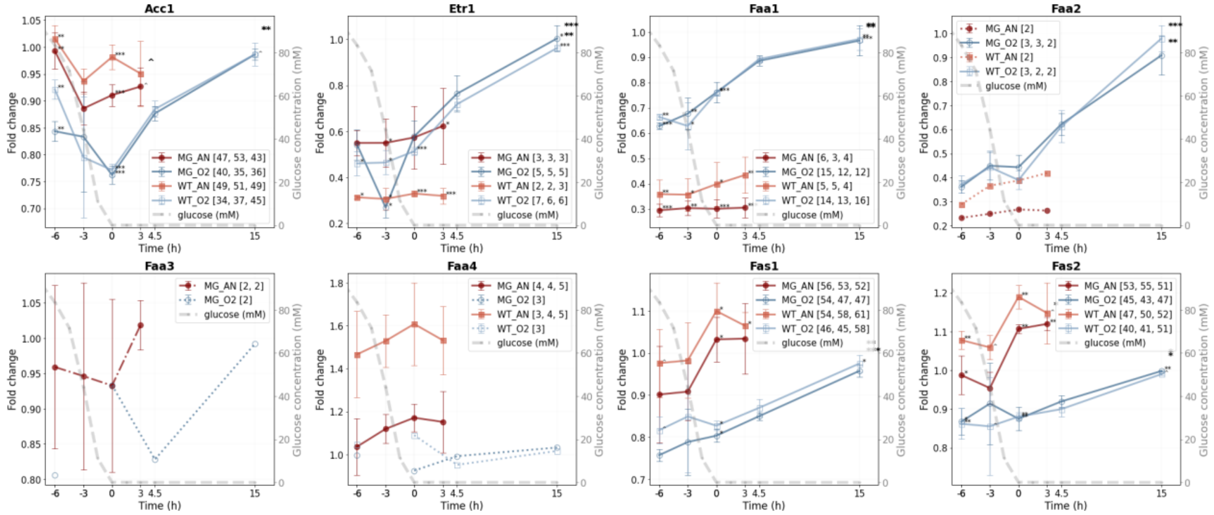


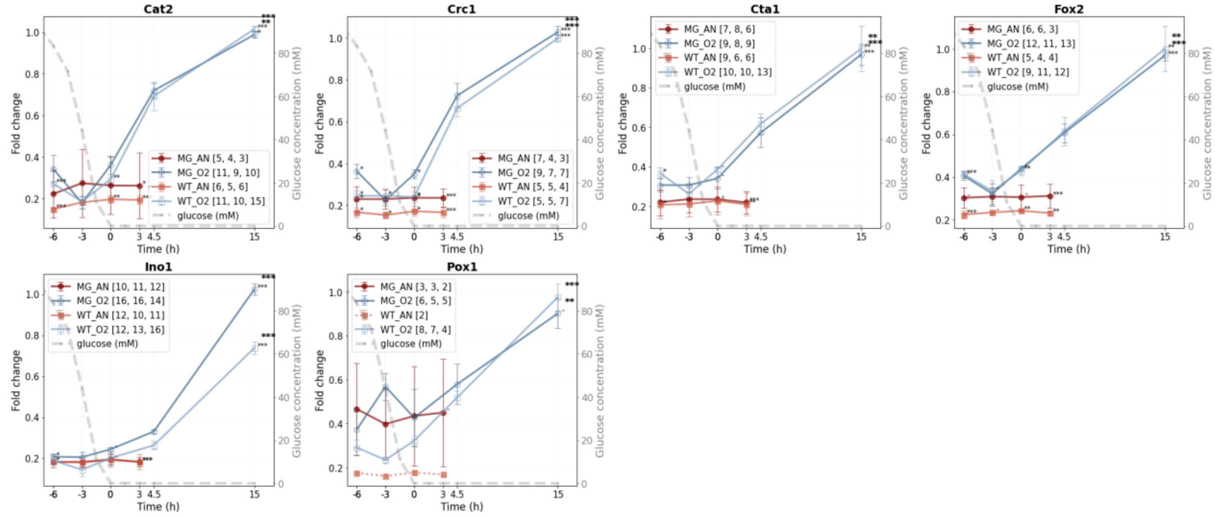

**SI Figure 6.4. Fold change line graphs for proteins of selected pathways for the control yeast and the MG strain under aerobic and anaerobic conditions.** The biological-replicate-averaged FCs of CEN.PK113-7D and IMX372 (MG) were plotted against the time relative to glucose depletion (0 h) in hours. Red: the MG strain under anaerobic conditions, dark blue: the MG strain under aerobic conditions, orange: the control strain under anaerobic conditions, light blue: the control strain under aerobic conditions. The error bars show the standard deviation of the mean of the three biological replicates. In the legend, the numbers between brackets represent the number of unique peptides that were found in each biological replicate. The grey dashed line represents the glucose concentration over time (mM, secondary y-axis). Asterisks (*) and circumflexes (ˆ) indicate the significance between either the (an)aerobic experiments (black annotation) or between the ME and MS phase. P-value levels are: $p < 0.001$ (***), $p < 0.01$ (**), $p < 0.05$ (*), and $p < 0.1$ (ˆ).

### Autophagy

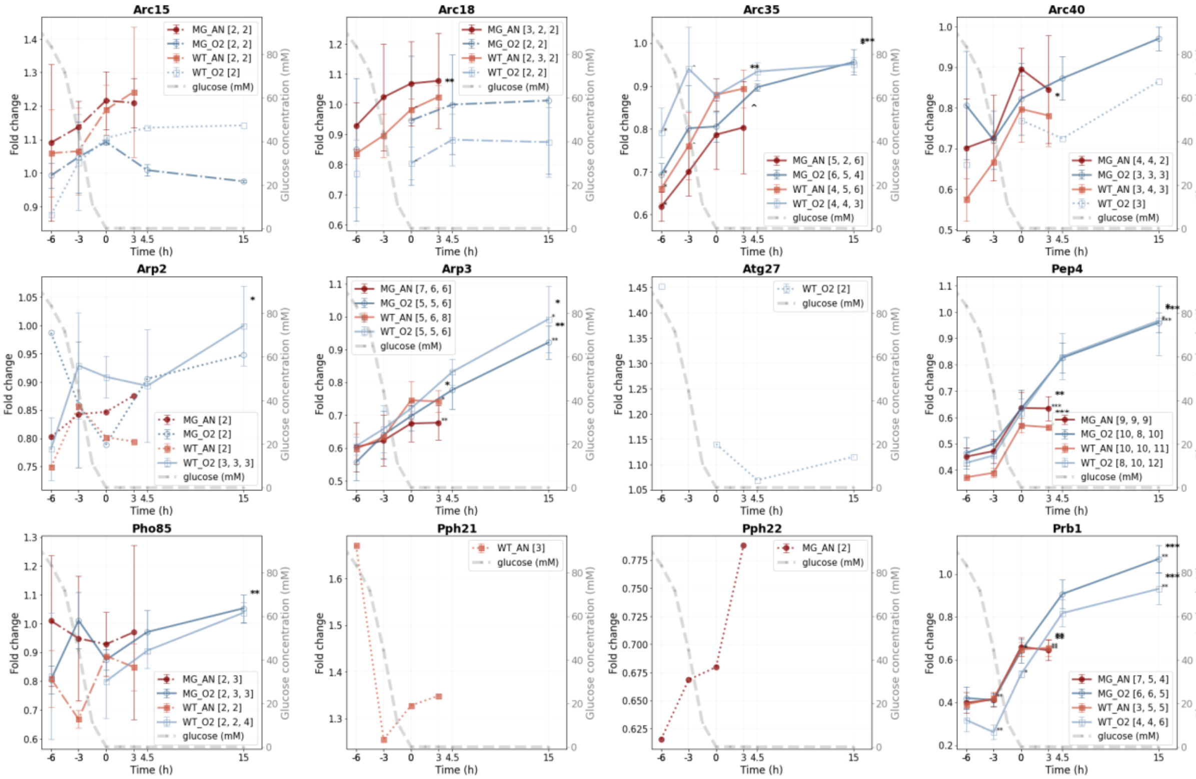
Arc15
MG_AN [2, 2]
MG_O2 [2, 2]
WT_AN [2, 2]
WT_O2 [2]
glucose (mM)
Arc18
MG_AN [3, 2, 2]
MG_O2 [2, 2]
WT_AN [2, 3, 2]
WT_O2 [2, 2]
glucose (mM)
Arc35
MG_AN [5, 2, 6]
MG_O2 [6, 5, 4]
WT_AN [4, 5, 6]
WT_O2 [4, 4, 3]
glucose (mM)
Arc40
MG_AN [4, 4, 2]
MG_O2 [3, 3, 3]
WT_AN [3, 4, 3]
WT_O2 [3]
glucose (mM)
Arp2
MG_AN [2]
MG_O2 [2]
WT_AN [2]
WT_O2 [3, 3, 3]
glucose (mM)
Arp3
MG_AN [7, 6, 6]
MG_O2 [5, 5, 6]
WT_AN [5, 6, 8]
WT_O2 [5, 5, 6]
glucose (mM)
Atg27
WT_O2 [2]
glucose (mM)
Pep4
MG_AN [9, 9, 9]
MG_O2 [10, 8, 10]
WT_AN [10, 10, 11]
WT_O2 [8, 10, 12]
glucose (mM)
Pho85
MG_AN [2, 3]
MG_O2 [2, 3, 3]
WT_AN [2, 2]
WT_O2 [2, 2, 4]
glucose (mM)
Pph21
WT_AN [3]
glucose (mM)
Pph22
MG_AN [2]
glucose (mM)
Prb1
MG_AN [7, 5, 4]
MG_O2 [6, 6, 5]
WT_AN [3, 5, 5]
WT_O2 [4, 4, 6]
glucose (mM)
Fold change
Glucose concentration (mM)
Time (h)

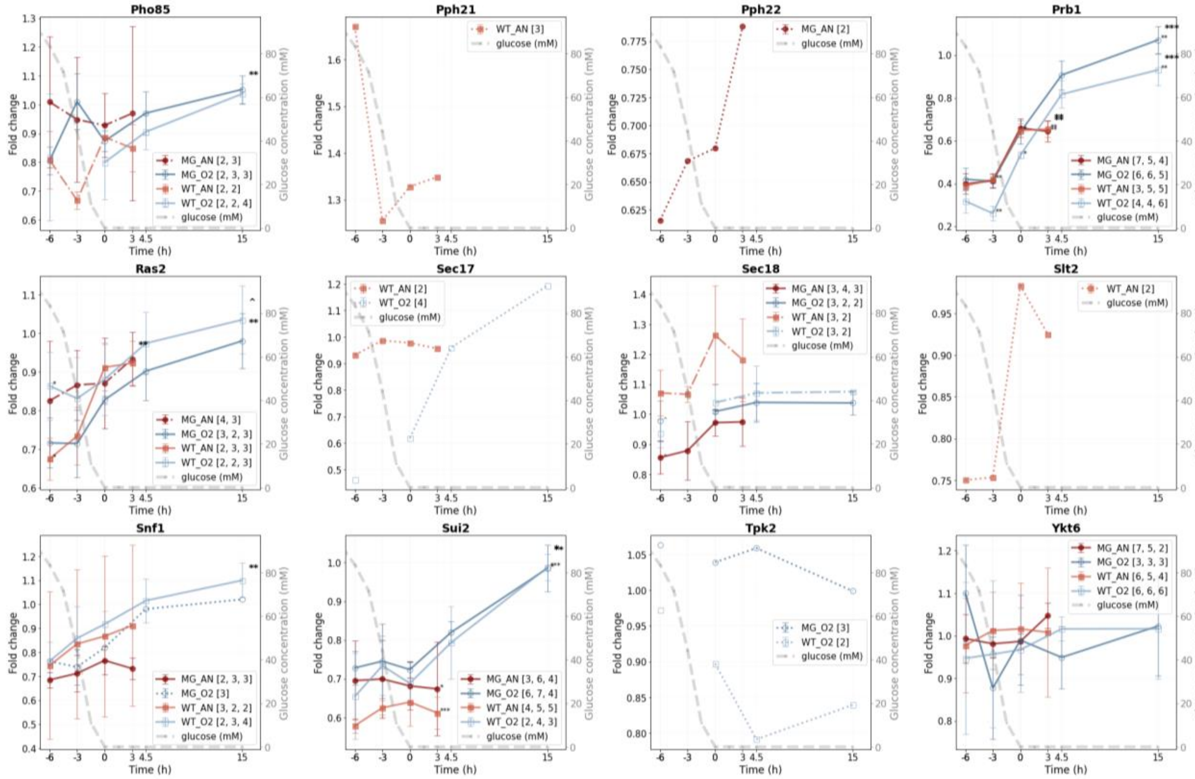
Pho85
MG_AN [2, 3]
MG_O2 [2, 3, 3]
WT_AN [2, 2]
WT_O2 [2, 2, 4]
glucose (mM)
Pph21
WT_AN [3]
glucose (mM)
Pph22
MG_AN [2]
glucose (mM)
Prb1
MG_AN [7, 5, 4]
MG_O2 [6, 6, 5]
WT_AN [3, 5, 5]
WT_O2 [4, 4, 6]
glucose (mM)
Ras2
MG_AN [4, 3]
MG_O2 [3, 2, 3]
WT_AN [2, 3, 3]
WT_O2 [2, 2, 3]
glucose (mM)
Sec17
WT_AN [2]
WT_O2 [4]
glucose (mM)
Sec18
MG_AN [3, 4, 3]
MG_O2 [3, 2, 2]
WT_AN [3, 2]
WT_O2 [3, 2]
glucose (mM)
Slt2
WT_AN [2]
glucose (mM)
Snf1
MG_AN [2, 3, 3]
MG_O2 [3]
WT_AN [3, 2, 2]
WT_O2 [2, 3, 4]
glucose (mM)
Sui2
MG_AN [3, 6, 4]
MG_O2 [6, 7, 4]
WT_AN [4, 5, 5]
WT_O2 [2, 4, 3]
glucose (mM)
Tpk2
MG_O2 [3]
WT_O2 [2]
glucose (mM)
Ykt6
MG_AN [7, 5, 2]
MG_O2 [3, 3, 3]
WT_AN [6, 5, 4]
WT_O2 [6, 6, 6]
glucose (mM)
Fold change
Glucose concentration (mM)
Time (h)

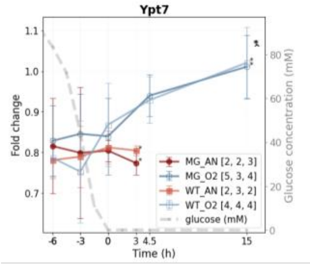


**SI Figure 6.5. Fold change line graphs for proteins of selected pathways for the control yeast and the MG strain under aerobic and anaerobic conditions.** The biological-replicate-averaged FCs of CEN.PK113-7D and IMX372 (MG) were plotted against the time relative to glucose depletion (0 h) in hours. Red: the MG strain under anaerobic conditions, dark blue: the MG strain under aerobic conditions, orange: the control strain under anaerobic conditions, light blue: the control strain under aerobic conditions. The error bars show the standard deviation of the mean of the three biological replicates. In the legend, the numbers between brackets represent the number of unique peptides that were found in each biological replicate. The grey dashed line represents the glucose concentration over time (mM, secondary y-axis). Asterisks (*) and circumflexes (ˆ) indicate the significance between either the (an)aerobic experiments (black annotation) or between the ME and MS phase. P-value levels are: p < 0.001 (***), p < 0.01 (**), p < 0.05 (*), and p < 0.1 (ˆ).

### Ribosomes

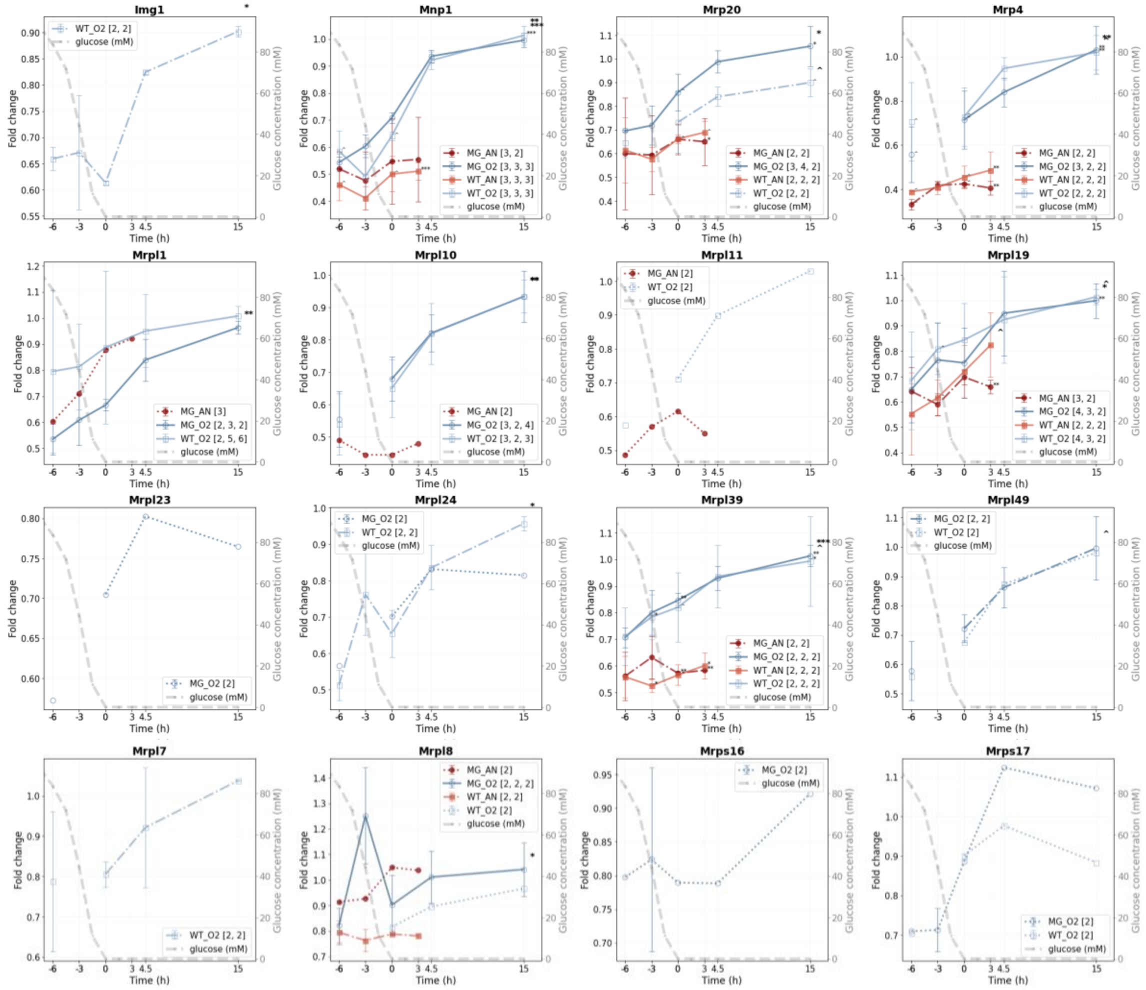

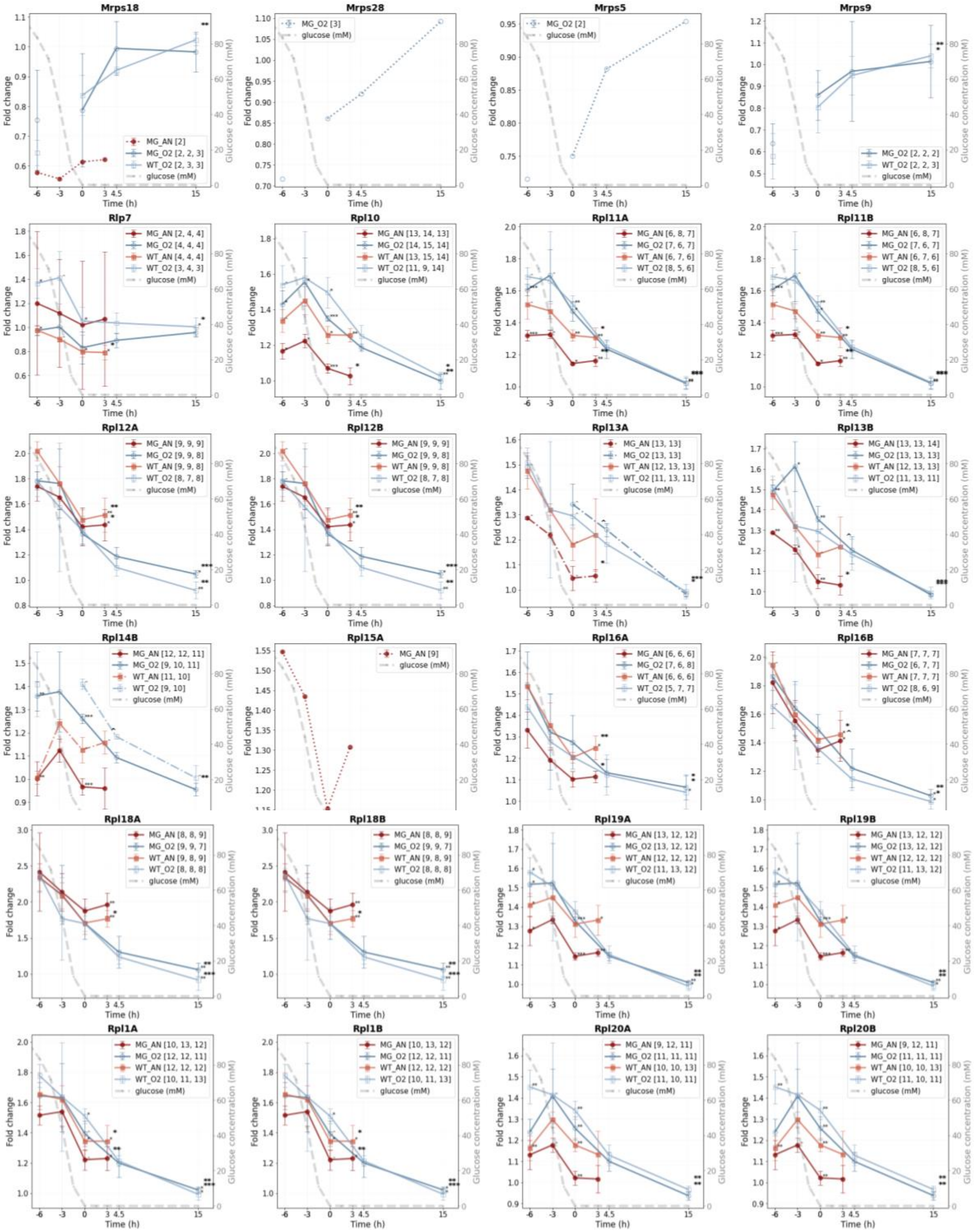
Mrps18
Mrps28
Mrps5
Mrps9
Rlp7
Rpl10
Rpl11A
Rpl11B
Rpl12A
Rpl12B
Rpl13A
Rpl13B
Rpl14B
Rpl15A
Rpl16A
Rpl16B
Rpl18A
Rpl18B
Rpl19A
Rpl19B
Rpl1A
Rpl1B
Rpl20A
Rpl20B
Fold change
Glucose concentration (mM)
Time (h)
glucose (mM)

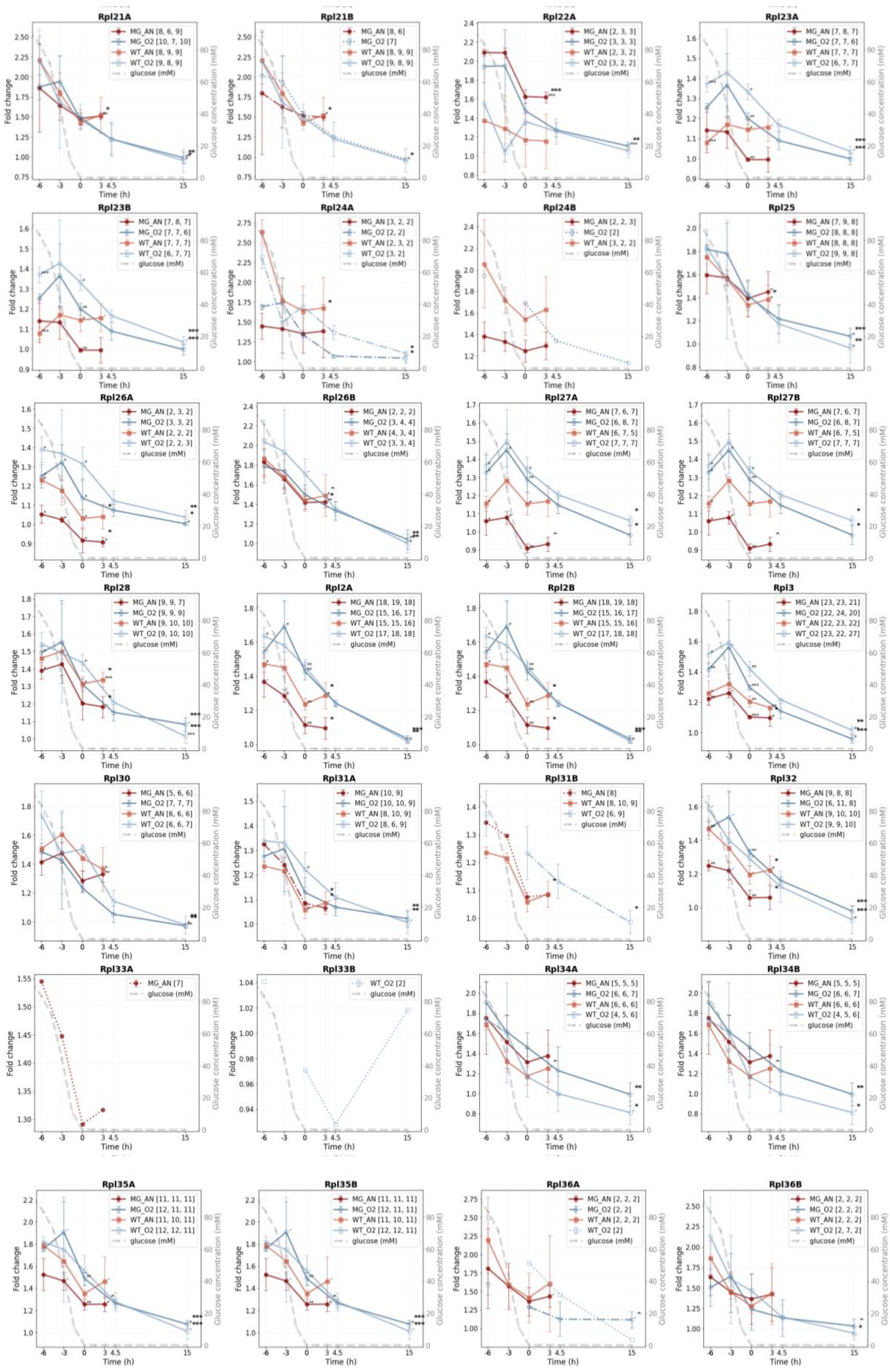
Rpl21A
MG_AN [8, 6, 9]
MG_O2 [10, 7, 10]
WT_AN [8, 9, 9]
WT_O2 [9, 8, 9]
glucose (mM)
Rpl21B
MG_AN [8, 6]
MG_O2 [7]
WT_AN [8, 9, 9]
WT_O2 [9, 8, 9]
glucose (mM)
Rpl22A
MG_AN [2, 3, 3]
MG_O2 [3, 3, 3]
WT_AN [2, 3, 2]
WT_O2 [3, 2, 2]
glucose (mM)
Rpl23A
MG_AN [7, 8, 7]
MG_O2 [7, 7, 6]
WT_AN [7, 7, 7]
WT_O2 [6, 7, 7]
glucose (mM)
Rpl23B
MG_AN [7, 8, 7]
MG_O2 [7, 7, 6]
WT_AN [7, 7, 7]
WT_O2 [6, 7, 7]
glucose (mM)
Rpl24A
MG_AN [3, 2, 2]
MG_O2 [2, 2]
WT_AN [2, 3, 2]
WT_O2 [3, 2]
glucose (mM)
Rpl24B
MG_AN [2, 2, 3]
MG_O2 [2]
WT_AN [3, 2, 2]
glucose (mM)
Rpl25
MG_AN [7, 9, 8]
MG_O2 [8, 8, 8]
WT_AN [8, 8, 8]
WT_O2 [9, 9, 8]
glucose (mM)
Rpl26A
MG_AN [2, 3, 2]
MG_O2 [3, 3, 2]
WT_AN [2, 2, 2]
WT_O2 [2, 2, 3]
glucose (mM)
Rpl26B
MG_AN [2, 2, 2]
MG_O2 [3, 4, 4]
WT_AN [4, 3, 4]
WT_O2 [3, 3, 4]
glucose (mM)
Rpl27A
MG_AN [7, 6, 7]
MG_O2 [6, 8, 7]
WT_AN [6, 7, 5]
WT_O2 [7, 7, 7]
glucose (mM)
Rpl27B
MG_AN [7, 6, 7]
MG_O2 [6, 8, 7]
WT_AN [6, 7, 5]
WT_O2 [7, 7, 7]
glucose (mM)
Rpl28
MG_AN [9, 9, 7]
MG_O2 [9, 9, 9]
WT_AN [9, 10, 10]
WT_O2 [9, 10, 10]
glucose (mM)
Rpl2A
MG_AN [18, 19, 18]
MG_O2 [15, 16, 17]
WT_AN [15, 15, 16]
WT_O2 [17, 18, 18]
glucose (mM)
Rpl2B
MG_AN [18, 19, 18]
MG_O2 [15, 16, 17]
WT_AN [15, 15, 16]
WT_O2 [17, 18, 18]
glucose (mM)
Rpl3
MG_AN [23, 23, 21]
MG_O2 [22, 24, 20]
WT_AN [22, 23, 22]
WT_O2 [23, 22, 27]
glucose (mM)
Rpl30
MG_AN [5, 6, 6]
MG_O2 [7, 7, 7]
WT_AN [6, 6, 6]
WT_O2 [6, 6, 7]
glucose (mM)
Rpl31A
MG_AN [10, 9]
MG_O2 [10, 10, 9]
WT_AN [8, 10, 9]
WT_O2 [8, 6, 9]
glucose (mM)
Rpl31B
MG_AN [8]
WT_AN [8, 10, 9]
WT_O2 [6, 9]
glucose (mM)
Rpl32
MG_AN [9, 8, 8]
MG_O2 [6, 11, 8]
WT_AN [9, 10, 10]
WT_O2 [9, 9, 10]
glucose (mM)
Rpl33A
MG_AN [7]
glucose (mM)
Rpl33B
WT_O2 [2]
glucose (mM)
Rpl34A
MG_AN [5, 5, 5]
MG_O2 [6, 6, 7]
WT_AN [6, 6, 6]
WT_O2 [4, 5, 6]
glucose (mM)
Rpl34B
MG_AN [5, 5, 5]
MG_O2 [6, 6, 7]
WT_AN [6, 6, 6]
WT_O2 [4, 5, 6]
glucose (mM)
Rpl35A
MG_AN [11, 11, 11]
MG_O2 [12, 11, 11]
WT_AN [11, 10, 11]
WT_O2 [12, 12, 11]
glucose (mM)
Rpl35B
MG_AN [11, 11, 11]
MG_O2 [12, 11, 11]
WT_AN [11, 10, 11]
WT_O2 [12, 12, 11]
glucose (mM)
Rpl36A
MG_AN [2, 2, 2]
MG_O2 [2, 2]
WT_AN [2, 2, 2]
WT_O2 [2]
glucose (mM)
Rpl36B
MG_AN [2, 2, 2]
MG_O2 [2, 2, 2]
WT_AN [2, 2, 2]
WT_O2 [2, 7, 2]
glucose (mM)
Fold change
Glucose concentration (mM)
Time (h)

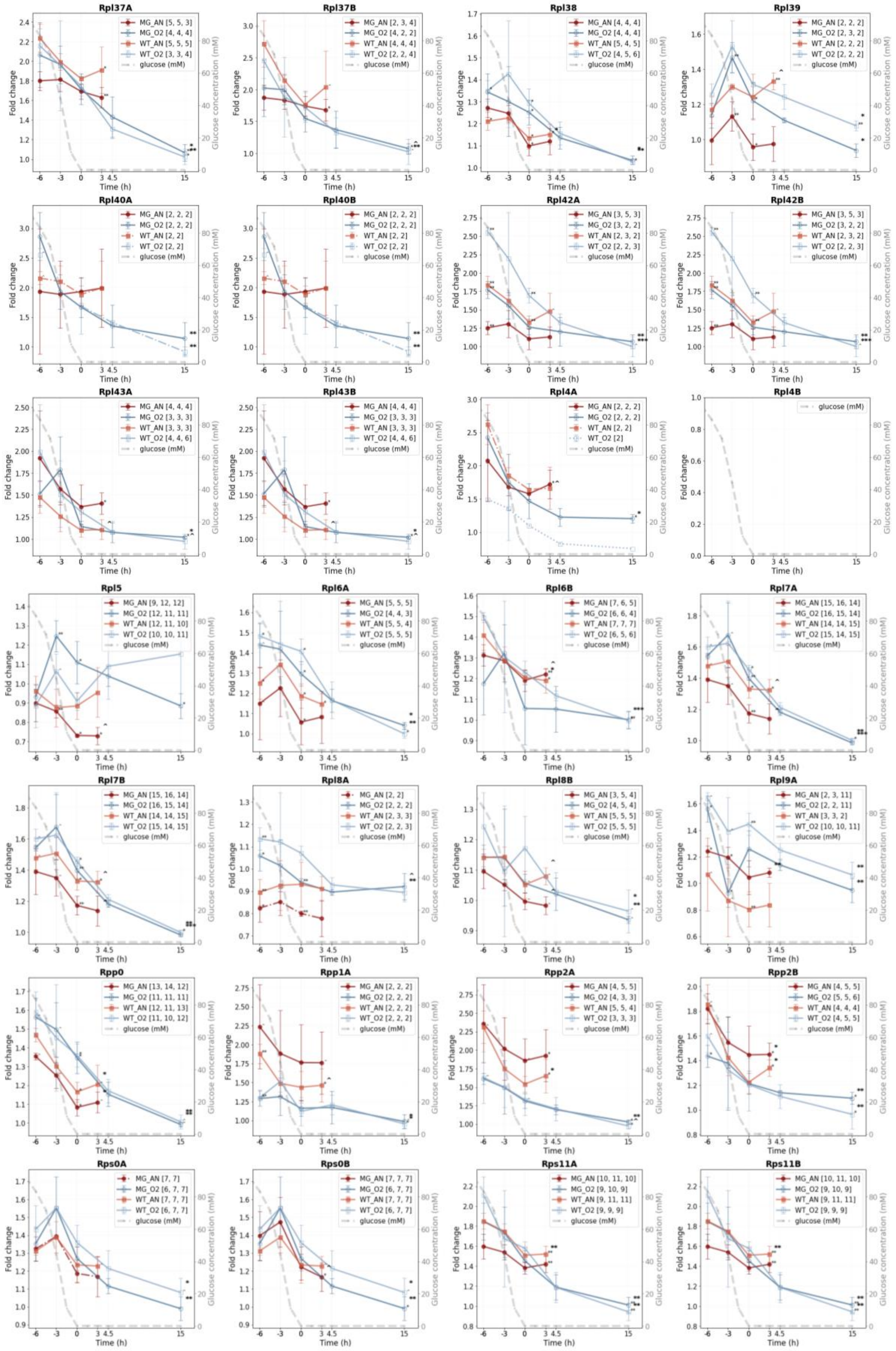
Rpl37A
MG_AN [5, 5, 3]
MG_O2 [4, 4, 4]
WT_AN [5, 5, 5]
WT_O2 [3, 3, 4]
glucose (mM)
Rpl37B
MG_AN [2, 3, 4]
MG_O2 [4, 2, 2]
WT_AN [4, 4, 4]
WT_O2 [2, 2, 4]
glucose (mM)
Rpl38
MG_AN [4, 4, 4]
MG_O2 [4, 4, 4]
WT_AN [5, 4, 5]
WT_O2 [4, 5, 6]
glucose (mM)
Rpl39
MG_AN [2, 2, 2]
MG_O2 [2, 3, 2]
WT_AN [2, 2, 2]
WT_O2 [2, 2, 2]
glucose (mM)
Rpl40A
MG_AN [2, 2, 2]
MG_O2 [2, 2, 2]
WT_AN [2, 2]
WT_O2 [2, 2]
glucose (mM)
Rpl40B
MG_AN [2, 2, 2]
MG_O2 [2, 2, 2]
WT_AN [2, 2]
WT_O2 [2, 2]
glucose (mM)
Rpl42A
MG_AN [3, 5, 3]
MG_O2 [3, 2, 2]
WT_AN [2, 3, 2]
WT_O2 [2, 2, 3]
glucose (mM)
Rpl42B
MG_AN [3, 5, 3]
MG_O2 [3, 2, 2]
WT_AN [2, 3, 2]
WT_O2 [2, 2, 3]
glucose (mM)
Rpl43A
MG_AN [4, 4, 4]
MG_O2 [3, 3, 3]
WT_AN [3, 3, 3]
WT_O2 [4, 4, 6]
glucose (mM)
Rpl43B
MG_AN [4, 4, 4]
MG_O2 [3, 3, 3]
WT_AN [3, 3, 3]
WT_O2 [4, 4, 6]
glucose (mM)
Rpl4A
MG_AN [2, 2, 2]
MG_O2 [2, 2, 2]
WT_AN [2, 2]
WT_O2 [2]
glucose (mM)
Rpl4B
glucose (mM)
Rpl5
MG_AN [9, 12, 12]
MG_O2 [12, 11, 11]
WT_AN [12, 11, 10]
WT_O2 [10, 10, 11]
glucose (mM)
Rpl6A
MG_AN [5, 5, 5]
MG_O2 [4, 4, 3]
WT_AN [5, 5, 4]
WT_O2 [5, 5, 5]
glucose (mM)
Rpl6B
MG_AN [7, 6, 5]
MG_O2 [6, 6, 4]
WT_AN [7, 7, 7]
WT_O2 [6, 5, 6]
glucose (mM)
Rpl7A
MG_AN [15, 16, 14]
MG_O2 [16, 15, 14]
WT_AN [14, 14, 15]
WT_O2 [15, 14, 15]
glucose (mM)
Rpl7B
MG_AN [15, 16, 14]
MG_O2 [16, 15, 14]
WT_AN [14, 14, 15]
WT_O2 [15, 14, 15]
glucose (mM)
Rpl8A
MG_AN [2, 2]
MG_O2 [2, 2, 2]
WT_AN [2, 3, 3]
WT_O2 [2, 2, 3]
glucose (mM)
Rpl8B
MG_AN [3, 5, 4]
MG_O2 [4, 5, 4]
WT_AN [5, 5, 5]
WT_O2 [5, 5, 5]
glucose (mM)
Rpl9A
MG_AN [2, 3, 11]
MG_O2 [2, 2, 11]
WT_AN [3, 3, 2]
WT_O2 [10, 10, 11]
glucose (mM)
Rpp0
MG_AN [13, 14, 12]
MG_O2 [11, 11, 11]
WT_AN [12, 11, 13]
WT_O2 [11, 10, 12]
glucose (mM)
Rpp1A
MG_AN [2, 2, 2]
MG_O2 [2, 2, 2]
WT_AN [2, 2, 2]
WT_O2 [2, 2, 2]
glucose (mM)
Rpp2A
MG_AN [4, 5, 5]
MG_O2 [4, 3, 3]
WT_AN [5, 5, 4]
WT_O2 [3, 3, 3]
glucose (mM)
Rpp2B
MG_AN [4, 5, 5]
MG_O2 [5, 5, 6]
WT_AN [4, 4, 4]
WT_O2 [4, 5, 5]
glucose (mM)
Rps0A
MG_AN [7, 7]
MG_O2 [6, 7, 7]
WT_AN [7, 7, 7]
WT_O2 [6, 7, 7]
glucose (mM)
Rps0B
MG_AN [7, 7, 7]
MG_O2 [6, 7, 7]
WT_AN [7, 7, 7]
WT_O2 [6, 7, 7]
glucose (mM)
Rps11A
MG_AN [10, 11, 10]
MG_O2 [9, 10, 9]
WT_AN [9, 11, 11]
WT_O2 [9, 9, 9]
glucose (mM)
Rps11B
MG_AN [10, 11, 10]
MG_O2 [9, 10, 9]
WT_AN [9, 11, 11]
WT_O2 [9, 9, 9]
glucose (mM)
Fold change
Glucose concentration (mM)
Time (h)

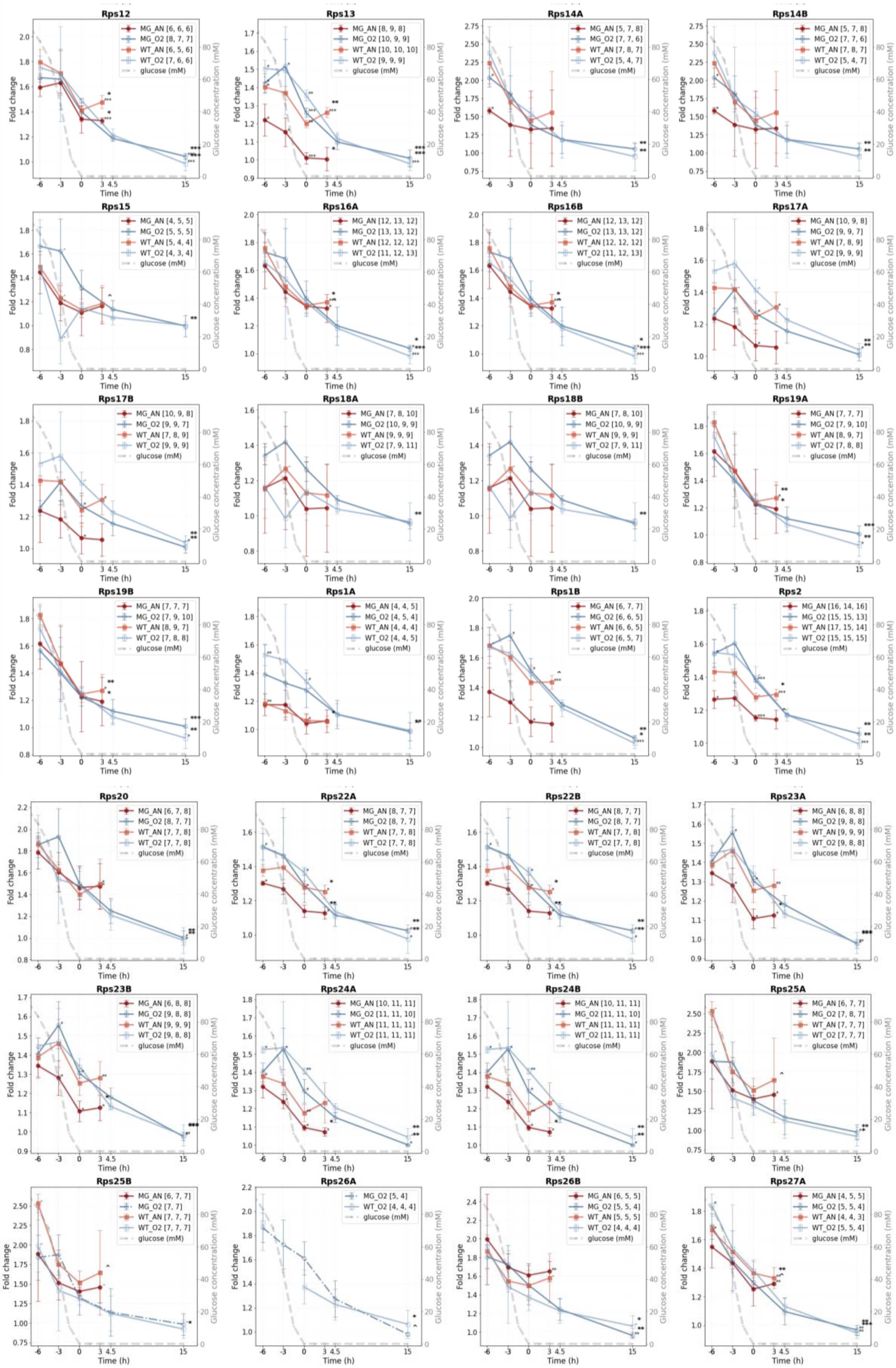
Rps12
Rps13
Rps14A
Rps14B
Rps15
Rps16A
Rps16B
Rps17A
Rps17B
Rps18A
Rps18B
Rps19A
Rps19B
Rps1A
Rps1B
Rps2
Rps20
Rps22A
Rps22B
Rps23A
Rps23B
Rps24A
Rps24B
Rps25A
Rps25B
Rps26A
Rps26B
Rps27A
Fold change
Glucose concentration (mM)
Time (h)
glucose (mM)

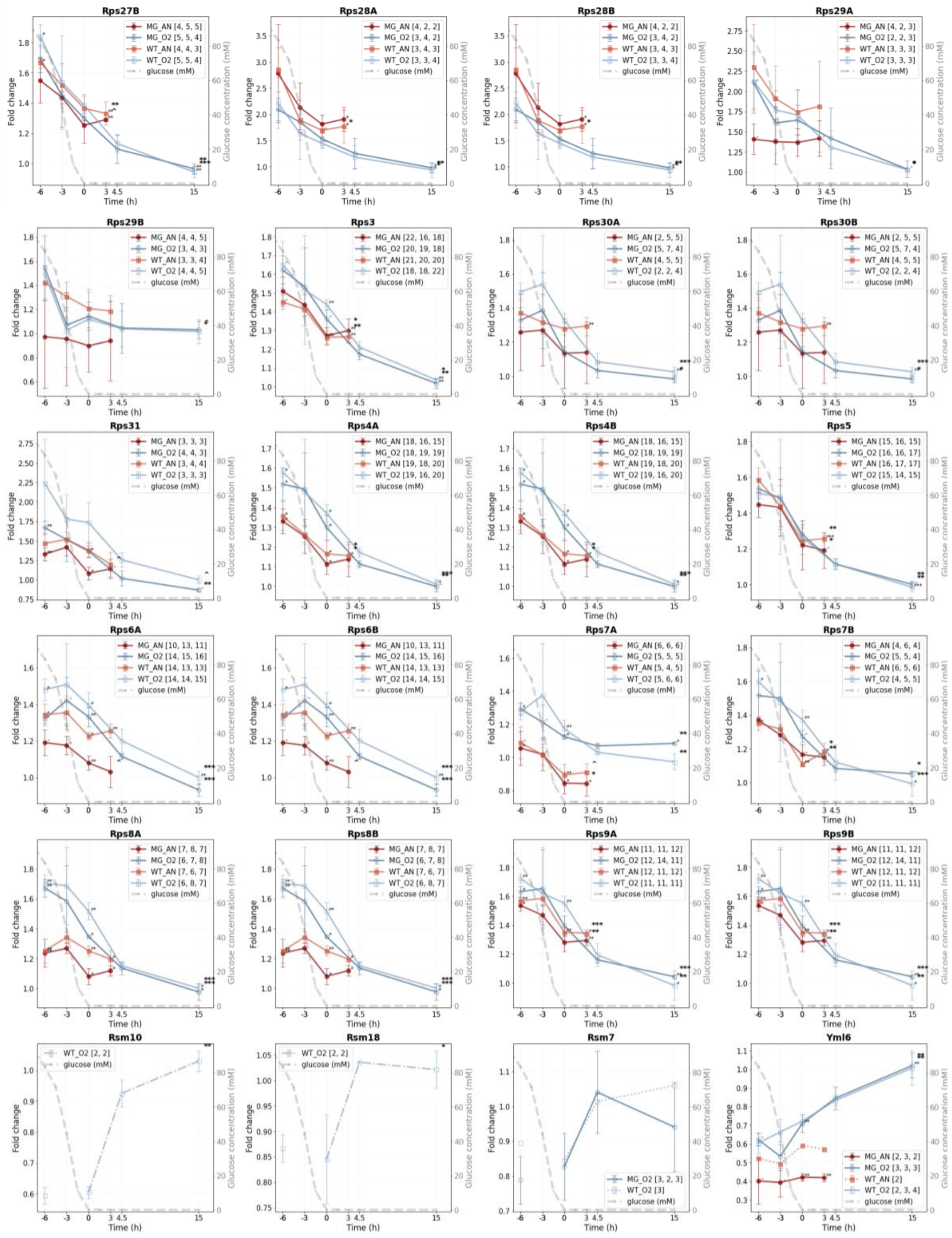


**SI Figure 6.6. Fold change line graphs for proteins of selected pathways for the control yeast and the MG strain under aerobic and anaerobic conditions.** The biological-replicate-averaged FCs of CEN.PK113-7D and IMX372 (MG) were plotted against the time relative to glucose depletion (0 h) in hours. Red: the MG strain under anaerobic conditions, dark blue: the MG strain under aerobic conditions, orange: the control strain under anaerobic conditions, light blue: the control strain under aerobic conditions. The error bars show the standard deviation of the mean of the three biological replicates. In the legend, the numbers between brackets represent the number of unique peptides that were found in each biological replicate. The grey dashed line represents the glucose concentration over time (mM, secondary y-axis). Asterisks (*) and circumflexes (^) indicate the significance between either the (an)aerobic experiments (black annotation) or between the ME and MS phase. P-value levels are: $p < 0.001$ (***), $p < 0.01$ (**), $p < 0.05$ (*), and $p < 0.1$ (^).

### ROS

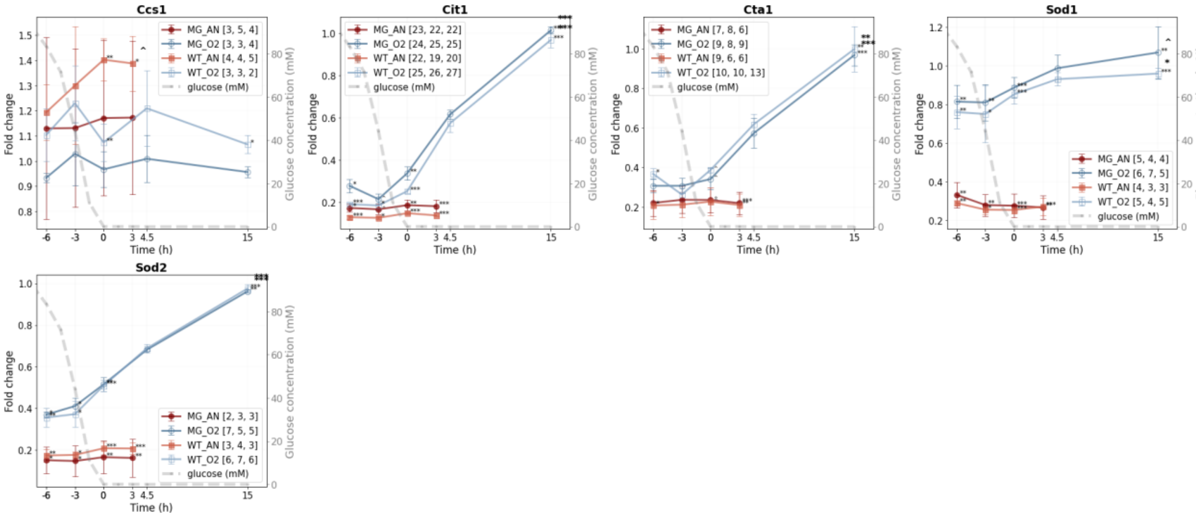


**SI Figure 6.7. Fold change line graphs for proteins of selected pathways for the control yeast and the MG strain under aerobic and anaerobic conditions.** The biological-replicate-averaged FCs of CEN.PK113-7D and IMX372 (MG) were plotted against the time relative to glucose depletion (0 h) in hours. Red: the MG strain under anaerobic conditions, dark blue: the MG strain under aerobic conditions, orange: the control strain under anaerobic conditions, light blue: the control strain under aerobic conditions. The error bars show the standard deviation of the mean of the three biological replicates. In the legend, the numbers between brackets represent the number of unique peptides that were found in each biological replicate. The grey dashed line represents the glucose concentration over time (mM, secondary y-axis). Asterisks (*) and circumflexes (^) indicate the significance between either the (an)aerobic experiments (black annotation) or between the ME and MS phase. P-value levels are: $p < 0.001$ (***), $p < 0.01$ (**), $p < 0.05$ (*), and $p < 0.1$ (^).

### Proteasome

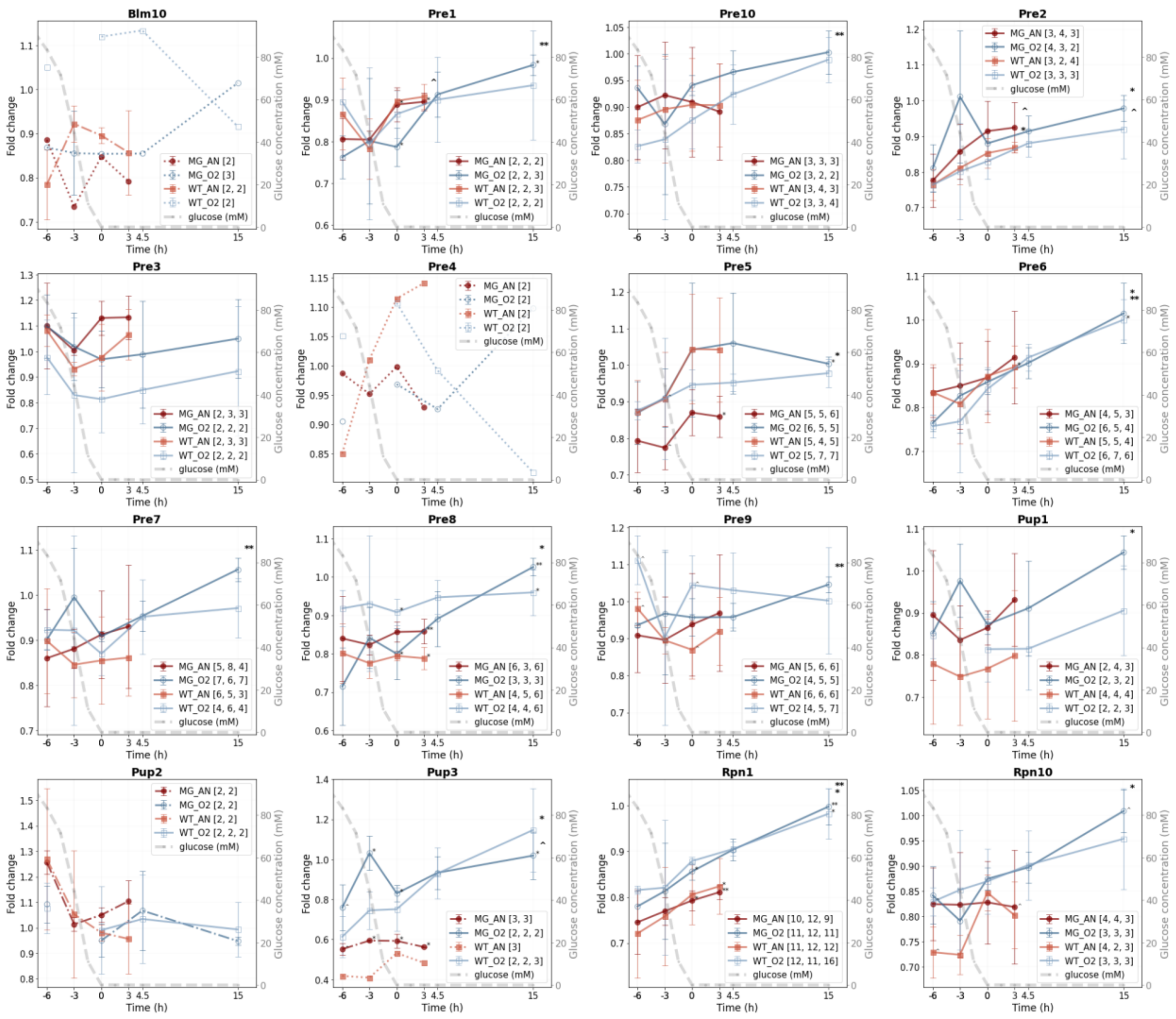
Blm10
MG_AN [2]
MG_O2 [3]
WT_AN [2, 2]
WT_O2 [2]
glucose (mM)
Pre1
MG_AN [2, 2, 2]
MG_O2 [2, 2, 3]
WT_AN [2, 2, 3]
WT_O2 [2, 2, 2]
glucose (mM)
Pre10
MG_AN [3, 3, 3]
MG_O2 [3, 2, 2]
WT_AN [3, 4, 3]
WT_O2 [3, 3, 4]
glucose (mM)
Pre2
MG_AN [3, 4, 3]
MG_O2 [4, 3, 2]
WT_AN [3, 2, 4]
WT_O2 [3, 3, 3]
glucose (mM)
Pre3
MG_AN [2, 3, 3]
MG_O2 [2, 2, 2]
WT_AN [2, 3, 3]
WT_O2 [2, 2, 2]
glucose (mM)
Pre4
MG_AN [2]
MG_O2 [2]
WT_AN [2]
WT_O2 [2]
glucose (mM)
Pre5
MG_AN [5, 5, 6]
MG_O2 [6, 5, 5]
WT_AN [5, 4, 5]
WT_O2 [5, 7, 7]
glucose (mM)
Pre6
MG_AN [4, 5, 3]
MG_O2 [6, 5, 4]
WT_AN [5, 5, 4]
WT_O2 [6, 7, 6]
glucose (mM)
Pre7
MG_AN [5, 8, 4]
MG_O2 [7, 6, 7]
WT_AN [6, 5, 3]
WT_O2 [4, 6, 4]
glucose (mM)
Pre8
MG_AN [6, 3, 6]
MG_O2 [3, 3, 3]
WT_AN [4, 5, 6]
WT_O2 [4, 4, 6]
glucose (mM)
Pre9
MG_AN [5, 6, 6]
MG_O2 [4, 5, 5]
WT_AN [6, 6, 6]
WT_O2 [4, 5, 7]
glucose (mM)
Pup1
MG_AN [2, 4, 3]
MG_O2 [2, 3, 2]
WT_AN [4, 4, 4]
WT_O2 [2, 2, 3]
glucose (mM)
Pup2
MG_AN [2, 2]
MG_O2 [2, 2]
WT_AN [2, 2]
WT_O2 [2, 2, 2]
glucose (mM)
Pup3
MG_AN [3, 3]
MG_O2 [2, 2, 2]
WT_AN [3]
WT_O2 [2, 2, 3]
glucose (mM)
Rpn1
MG_AN [10, 12, 9]
MG_O2 [11, 12, 11]
WT_AN [11, 12, 12]
WT_O2 [12, 11, 16]
glucose (mM)
Rpn10
MG_AN [4, 4, 3]
MG_O2 [3, 3, 3]
WT_AN [4, 2, 3]
WT_O2 [3, 3, 3]
glucose (mM)
Fold change
Glucose concentration (mM)
Time (h)

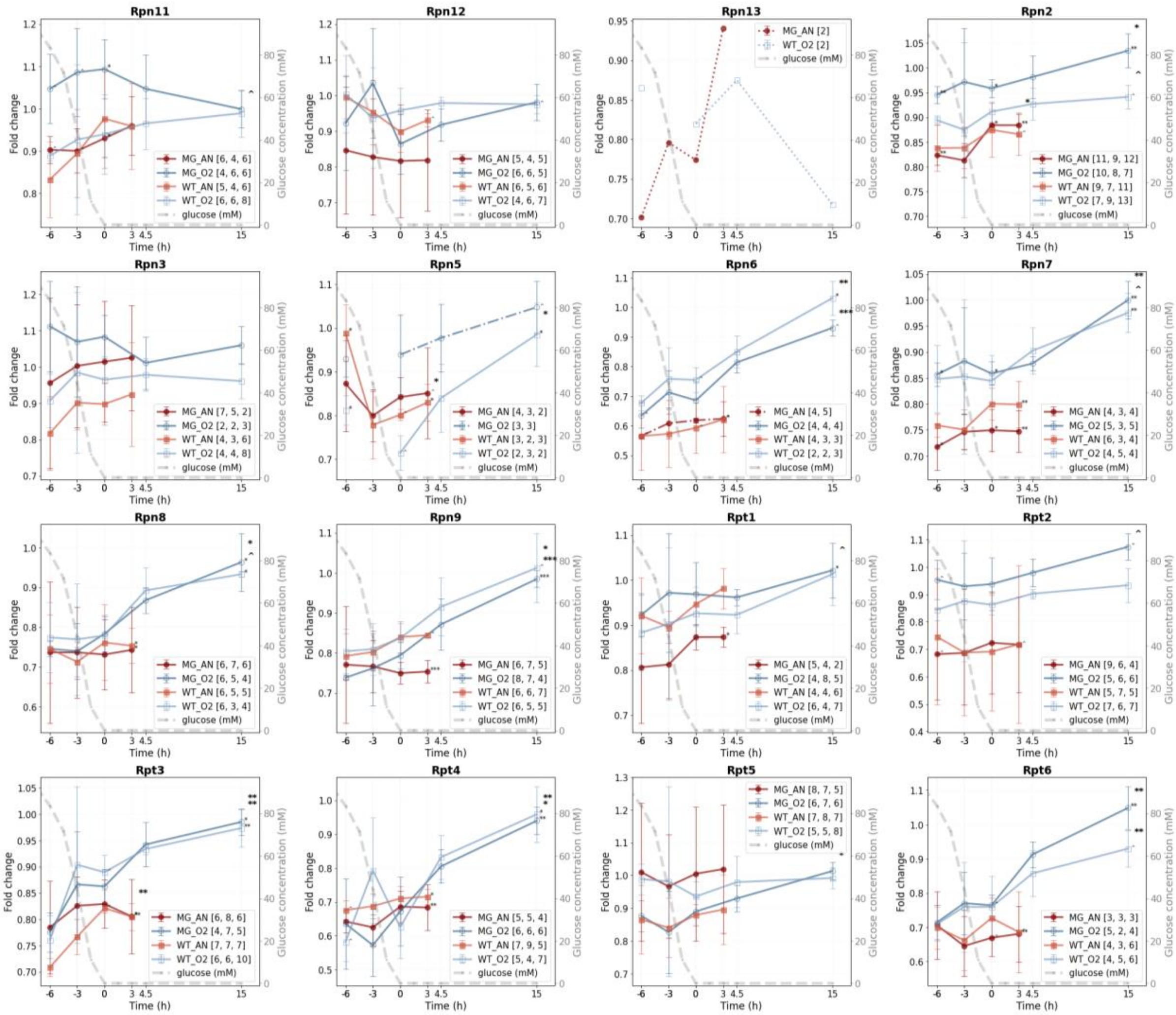


**SI Figure 6.8. Fold change line graphs for proteins of selected pathways for the control yeast and the MG strain under aerobic and anaerobic conditions.** The biological-replicate-averaged FCs of CEN.PK113-7D and IMX372 (MG) were plotted against the time relative to glucose depletion (0 h) in hours. Red: the MG strain under anaerobic conditions, dark blue: the MG strain under aerobic conditions, orange: the control strain under anaerobic conditions, light blue: the control strain under aerobic conditions. The error bars show the standard deviation of the mean of the three biological replicates. In the legend, the numbers between brackets represent the number of unique peptides that were found in each biological replicate. The grey dashed line represents the glucose concentration over time (mM, secondary y-axis). Asterisks (*) and circumflexes (ˆ) indicate the significance between either the (an)aerobic experiments (black annotation) or between the ME and MS phase. P-value levels are: $p < 0.001$ (***), $p < 0.01$ (**), $p < 0.05$ (*), and $p < 0.1$ (ˆ).

### Protein degradation 26S proteasome complex

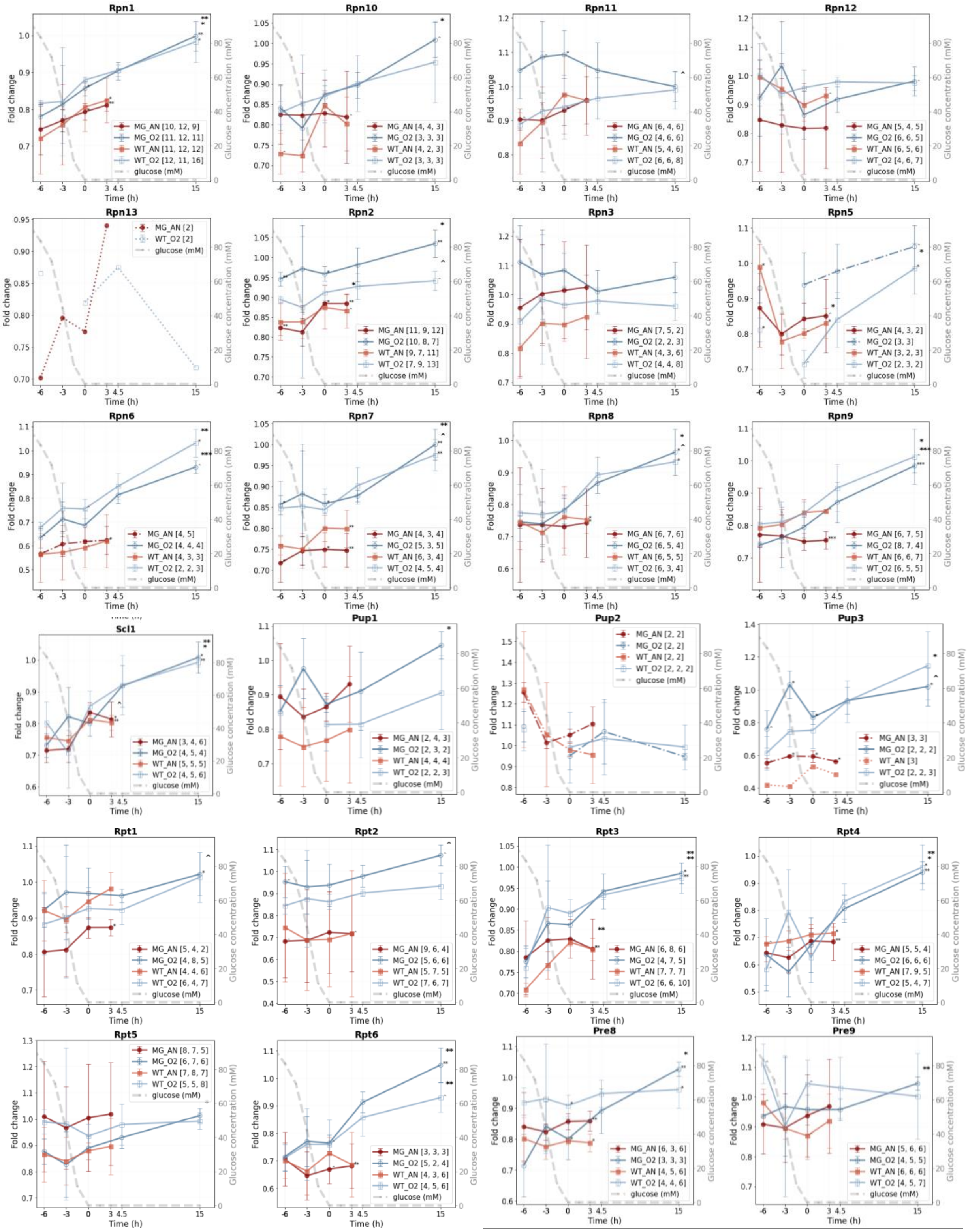

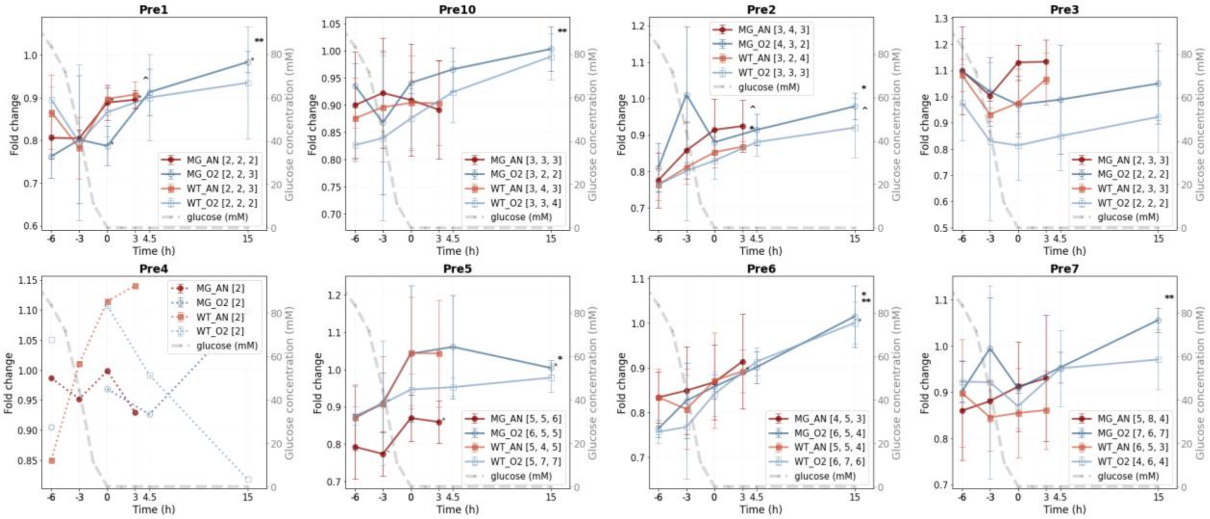


**SI Figure 6.9. Fold change line graphs for proteins of selected pathways for the control yeast and the MG strain under aerobic and anaerobic conditions.** The biological-replicate-averaged FCs of CEN.PK113-7D and IMX372 (MG) were plotted against the time relative to glucose depletion (0 h) in hours. Red: the MG strain under anaerobic conditions, dark blue: the MG strain under aerobic conditions, orange: the control strain under anaerobic conditions, light blue: the control strain under aerobic conditions. The error bars show the standard deviation of the mean of the three biological replicates. In the legend, the numbers between brackets represent the number of unique peptides that were found in each biological replicate. The grey dashed line represents the glucose concentration over time (mM, secondary y-axis). Asterisks (*) and circumflexes (ˆ) indicate the significance between either the (an)aerobic experiments (black annotation) or between the ME and MS phase. P-value levels are: $p < 0.001$ (***), $p < 0.01$ (**), $p < 0.05$ (*), and $p < 0.1$ (ˆ).

### Heme synthesis

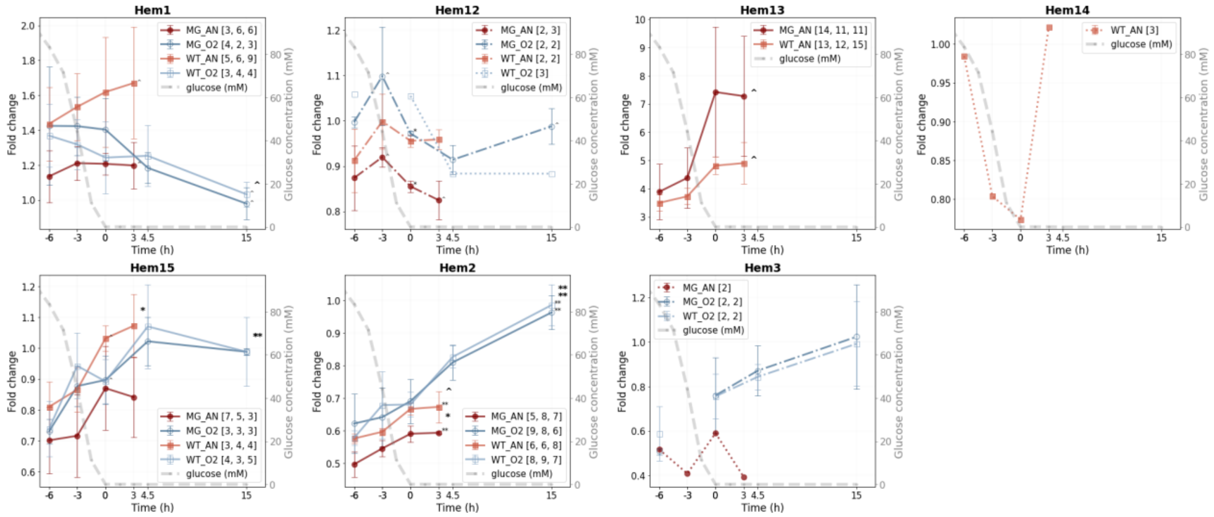

**SI Figure 6.10. Fold change line graphs for proteins of selected pathways for the control yeast and the MG strain under aerobic and anaerobic conditions.** The biological-replicate-averaged FCs of CEN.PK113-7D and IMX372 (MG) were plotted against the time relative to glucose depletion (0 h) in hours. Red: the MG strain under anaerobic conditions, dark blue: the MG strain under aerobic conditions, orange: the control strain under anaerobic conditions, light blue: the control strain under aerobic conditions. The error bars show the standard deviation of the mean of the three biological replicates. In the legend, the numbers between brackets represent the number of unique peptides that were found in each biological replicate. The grey dashed line represents the glucose concentration over time (mM, secondary y-axis). Asterisks (*) and circumflexes (ˆ) indicate the significance between either the (an)aerobic experiments (black annotation) or between the ME and MS phase. P-value levels are: $p < 0.001$ (***), $p < 0.01$ (**), $p < 0.05$ (*), and $p < 0.1$ (ˆ).

### Sterol synthesis

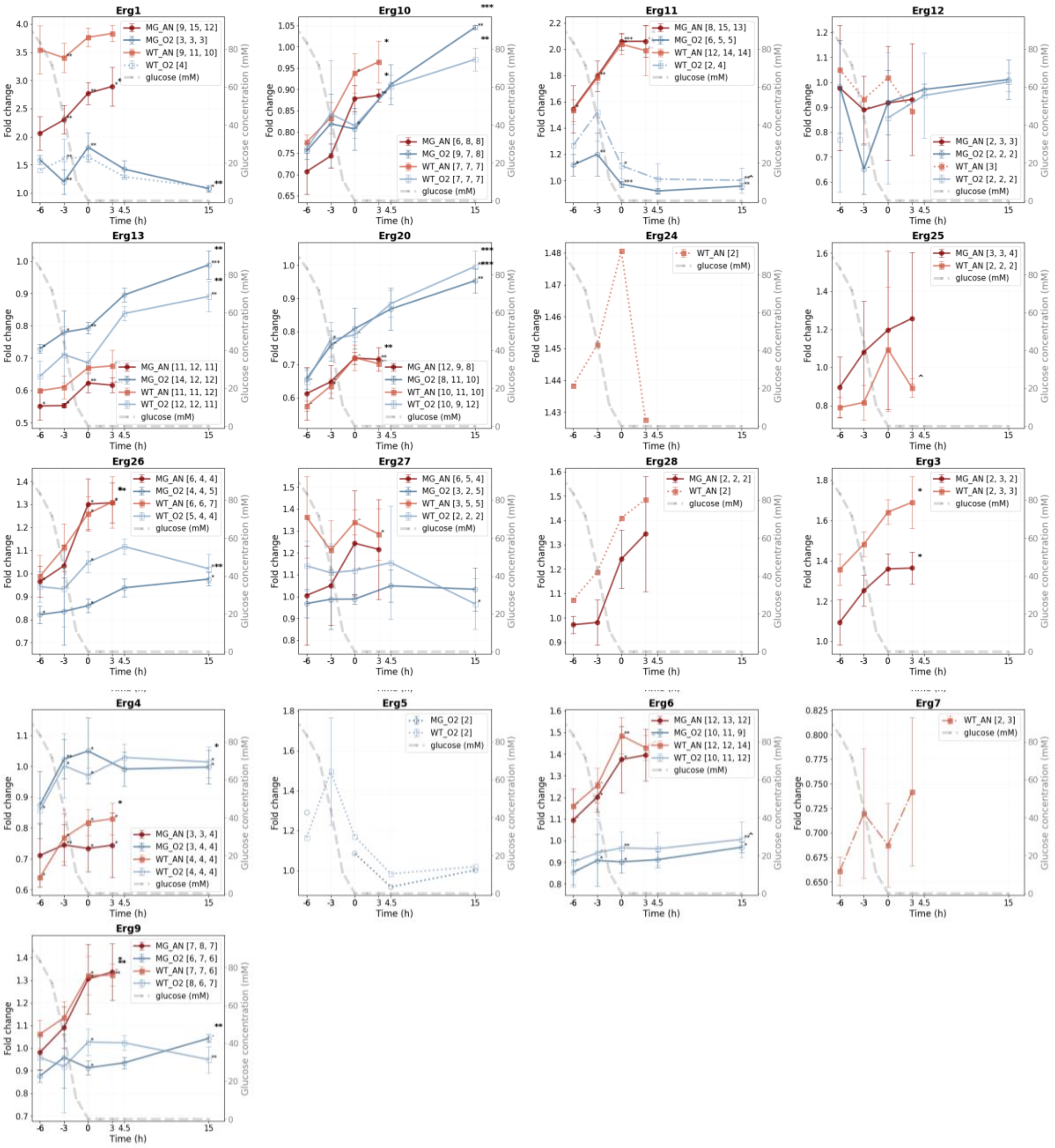


**SI Figure 6.11. Fold change line graphs for proteins of selected pathways for the control yeast and the MG strain under aerobic and anaerobic conditions.** The biological-replicate-averaged FCs of CEN.PK113-7D and IMX372 (MG) were plotted against the time relative to glucose depletion (0 h) in hours. Red: the MG strain under anaerobic conditions, dark blue: the MG strain under aerobic conditions, orange: the control strain under anaerobic conditions, light blue: the control strain under aerobic conditions. The error bars show the standard deviation of the mean of the three biological replicates. In the legend, the numbers between brackets represent the number of unique peptides that were found in each biological replicate. The grey dashed line represents the glucose concentration over time (mM, secondary y-axis). Asterisks (*) and circumflexes (ˆ) indicate the significance between either the (an)aerobic experiments (black annotation) or between the ME and MS phase. P-value levels are: $p < 0.001$ (***), $p < 0.01$ (**), $p < 0.05$ (*), and $p < 0.1$ (ˆ).
